# Biofilm inhibitor taurolithocholic acid alters colony morphology, specialized metabolism, and virulence of *Pseudomonas aeruginosa*

**DOI:** 10.1101/675405

**Authors:** Alanna R. Condren, Lisa Juliane Kahl, George Kritikos, Manuel Banzhaf, Lars E. P. Dietrich, Laura M. Sanchez

## Abstract

Biofilm inhibition by exogenous molecules has been an attractive strategy for the development of novel therapeutics. We investigated the biofilm inhibitor taurolithocholic acid (TLCA) and its effects on the specialized metabolism, virulence and biofilm formation of the clinically relevant bacterium *Pseudomonas aeruginosa* strain PA14. Our study shows that TLCA alters specialized metabolism, thereby affecting *P. aeruginosa* colony biofilm physiology. We observed an upregulation of metabolites correlated to virulence such as the siderophore pyochelin. A wax moth virulence assay confirmed that treatment with TLCA increases virulence of *P. aeruginosa*. Based on our results, we believe that future endeavors to identify biofilm inhibitors must consider how a putative lead is altering the specialized metabolism of a bacterial community to prevent pathogens from entering a highly virulent state.

## Introduction

The ESKAPE pathogens (*Enterococcus faecium, Staphylococcus aureus, Klebsiella pneumoniae, Acinetobacter baumannii, Pseudomonas aeruginosa,* and *Enterobacter* species) have been deemed a severe threat, as the major cause of nosocomial infections, by evolving mechanisms to “escape” the biocidal action of antibiotics [1,2]. The Center for Disease Control and Prevention estimates that costs related to nosocomial infections, which have increased in frequency in all countries regardless of income or industrial development, are between $680 to $5,683 USD on average per patient [3].

*P. aeruginosa,* one of the ESKAPE microorganisms, is often referred as a “ubiquitous” bacterium because of its ability to adapt to a wide variety of environments and hosts [4]. *P. aeruginosa* can be found in the lung cavity of cystic fibrosis patients, colonizing large open wounds of burn victims in hospitals, or invading the cornea of the human eye leading to permanent vision loss [5–7]. *P. aeruginosa* infections are often complicated by the fact that it readily forms a multicellular aggregate known as a biofilm - a state which contributes towards its resistance to antibiotics [8]. The minimum bactericidal concentration for cells in a biofilm state is estimated to be 10-1000 times higher than their planktonic counterparts complicating treatment of biofilm infections [9]. *P. aeruginosa* tightly regulates biofilm formation using molecular signaling networks and a well-characterized arsenal of specialized metabolites [10–14]. Specialized metabolites encompass the bioactive small molecules produced by a given organism outside of those needed for primary metabolism [15]. Yet, there are currently no biofilm inhibitors on the market in the US [16].

There are, however, several biofilm inhibitors used for *in vitro* analysis of biofilm dispersal [17]. One example is taurolithocholic acid (TLCA), a bile acid which efficiently inhibits biofilm formation and induces dispersion of mature *P. aeruginosa* biofilms [18,19]. TLCA demonstrated a low micromolar biofilm inhibitory concentration (BIC_50_) against *P. aeruginosa* at 38.4 μM compared to other lithocholic and bile acid derivatives [18]. Bile acids are a class of acidic steroids that play a physiological role in digestion by solubilizing dietary fats [20]. Though bile acids are classified as detergents, the steroid control cholesterol 3-sulfate, does not inhibit biofilm formation nor do other bile acids tested at concentrations up to 1 mM. However, all lithocholic bile acids have specific bioactivity against *P. aeruginosa* which varies based on the conjugation to glycine or taurine [18]. Additionally, the reported BIC_50_ of TLCA was in the low micromolar range while the maximum critical micelle concentration for these bile acids ranges from 8 to 12 mM [20]. Therefore, we hypothesized that when TLCA enters the microorganism’s chemical space, it induces *P. aeruginosa* to alter the production of specific specialized metabolites leading to the reported biofilm inhibition/dispersion.

We observed altered morphology of colony biofilms and changes in specialized metabolism when *P. aeruginosa* PA14 is exposed to TLCA. An increase in pyochelin production when TLCA is present lead us to perform a wax moth virulence model which confirmed that TLCA treated cells are significantly more virulent than non-treated cells [21]. The observed increase in virulence lead to an investigation into how TLCA may be inducing an increase in pyochelin production to increase virulence. We performed an iron starvation tolerance assay as well as mutant studies via imaging mass spectrometry (IMS) which concluded that TLCA treatment does not make *P. aeruginosa* cells sensitive to iron starvation and a knockout of the *pqsH* (Δ*pqsH*) and *phzA1-G1 phzA2-G2* (Δ*phz*) gene clusters also leads to an increase in pyochelin production as observed for the wild-type strain [22]. Taken together, while TLCA has shown promising bioactivity towards biofilm inhibition, it appears that biofilm inhibition (or dispersion) ultimately leads to the bacterium becoming more virulent in a host model, which is supported by the observed alteration in specialized metabolism.

## Methods & Materials

### TLCA stock solution preparation

Taurolithocholic acid was purchased from Sigma Aldrich (≥98%). A 0.5 M stock solution was made by dissolving TLCA in methanol. This solution was then sterile filtered with a 0.22 μm sterile filter and stored at −80□.

### Colony Morphology Studies

Liquid agar was prepared by autoclaving 1% tryptone (Teknova), 1% agar (Teknova) mixture. The liquid agar was cooled to 60°C and Congo Red (EMD; final concentration 40 μg/mL) and Coomassie Blue (EMD; final concentration: 20 μg/mL) were added. TLCA stock solution, dissolved in MeOH, was vortexed for 2-3 minutes until clear. TLCA was added to liquid agar, different amounts were added to reach the final concentrations of 100 μM, 250 μM, and 1 mM TLCA. Sixty mL of liquid agar mixture was poured into square plates (LDP, 10 cm × 10 cm × 1.5 cm) and left to solidify for ~18-24 hours. Precultures of *P. aeruginosa* PA14 were grown in LB for 12-16 h at 37°C while shaking at 250 rpm. Subcultures were prepared as 1:100 dilutions of precultures into fresh LB media and shaking for 2.5 hours at 37°C, at which point all subcultures had reached mid-exponential phase (optical density of ~0.4-0.6 at 500 nm). Morphology plates were dried for 20-30 min and 10 μl spots of subculture were spotted onto a morphology plate, with not more than four colonies per plate. Colony biofilms were grown at 25°C and high humidity (+90%) for up to 5 days. Images were taken every 24 hours with a Keyence VHX-1000 microscope.

### Imaging mass spectrometry experiments

TLCA and the respective vehicle were imbedded into liquid agar prior to plating. LB agar was autoclaved and cooled to 55 □ before adding TLCA. A calculated aliquot of sterilized TLCA stock solution was added to achieve the desired concentration of 250 μM. Plates were stored in 4□ refrigerator until needed. *P. aeruginosa* PA14 was plated on Bacto LB agar and grown overnight at 30□. A colony from the plate was then used to inoculate a 5 mL LB liquid culture of *P. aeruginosa* and grown overnight at 30□ shaking at 225 rpm. Overnight liquid cultures (5 μL) was spotted on thin agar plates (10 mL of agar in 90 mm plate) embedded with either TLCA or the vehicle (MeOH) and incubated for 48 hours at 30□. Humidity/moisture was removed from environment during growth by placing a small amount of DrieRite in the incubator. Following 48 hours of growth, colonies were excised from agar plates using a razor blade and transferred to an MSP 96 target ground steel target plate (Bruker Daltonics). An optical image of the colonies on the target plate was taken prior to matrix application. A 53 μm stainless steel sieve (Hogentogler Inc) was used to coat the steel target plate and colonies with MALDI matrix. The MALDI matrix used for the analysis was a 1:1 mixture of recrystallized □-cyano-4-hydroxycinnamic acid (CHCA) and 2,5-dihydroxybenzoic acid (DHB) (Sigma). The plate was then placed in an oven at 37□ for approximately four hours or until the agar was fully desiccated. After four hours, excess matrix was removed from target plate and sample with a stream of air. Another optical image was taken of the desiccated colonies on the target plate. The target plate and desiccated colony were then introduced into the MALDI-TOF mass spectrometer (Bruker Autoflex Speed) and analyzed with FlexControl v.3.4 and FlexImaging v.4.1 software. The detector gain and laser power must be optimized for each imaging run to achieve acceptable signal to noise, as an example, in figure 2, the detector gain and laser power were set at 3.0× and 41% respectively. Range of detection was from 100 Da to 3,500 Da with ion suppression set at 50 Da in positive reflectron mode. The raster size was set to 500 μm and the laser was set to 200 shots per raster point at medium (3) laser size.

### Statistical analysis of Imaging Data

SCiLS software (Bruker, version 2015b) was used to run statistical analysis of raw imaging data. Settings for analysis was as follows: Normalization: root mean square (RMS), error: ∓0.2 Da, and weak denoising for segmentation. Using “Find

Discriminative Values (ROC)” for unbiased analysis, PA14 control colony was selected as class 1 and TLCA treated colony as class 2 and SCiLS identified signals that were significantly upregulated in class 1 (MeOH control colony) with a threshold of 0.75 corresponding to a Pearson correlation of p<0.05. In our report, these signals are referred to as “downregulated” since they have a higher intensity in the control. The same analysis was performed with the classes flipped to identify signals that were upregulated in TLCA condition. Statistical analysis was completed after calculating mass error (**Table S2**) of three biological replicates (N=3) (**Figures S3-S5**) and signals were only considered significant if altered regulation was observed in two or more replicates.

### Iron Stress Tolerance Assay

The protocol for the iron stress tolerance assay used was exactly as described by Chua *et al.* [22], with the exceptions that strain PA14 rather than PAO1 and a 1 mM of TLCA and 2’2-bipyridine (DIPY) were used. The OD at 600 nm of each liquid culture was measured every 15 minutes for 16 hours.

### Galleria mellonella treatment assays

*Galleria mellonella* larvae (greater wax moth) were purchased from TrueLarv UK Ltd. (Exeter, UK) and stored at 15°C prior to use. The assay was performed as described previously [23], except for the following differences: PA14 wild-type (WT) cells were grown exponentially for 2 hours, washed with PBS buffer and adjusted to an OD_(600)_ of 0.1. The PA14 culture was further diluted with PBS and plated to determine the CFU of the inoculation suspension. Larvae were inoculated with 20 μL of a 2.5 × 10^3^ CFU/mL solution and incubated at 37°C for 2 hours. After this incubation time, larvae were injected with 20 μL of compound or PBS. The final concentration of TLCA and sodium nitroprusside (SNP) was 250 μM. For controls, uninfected larvae were administered the same compound dose to monitor for toxicity. In parallel one group of larvae received two sterile 20 μL PBS injections to control for unintentional killing by the injections. In total 25 larvae were used per condition, split in two independent experiments. In total 10 larvae were used for the PBS only condition. Survival of a larvae was determined by the ability to respond to external stimuli (poking). Larvae survival was estimated using the Kaplan-Meier estimator.[24] Survival estimates were subsequently compared using the log-rank test [25]. Resulting p-values were corrected using the Benjamini-Hochberg method [26] (**Figure S3**).

### pqs mutants IMS analysis

Mutants proceeded through the same protocol as the WT (PA14) bacterial colonies (**Table S10**). IMS sample prep and experimentation was completed at the same settings as described in “Imaging mass spectrometry experiments” section. Statistical analysis was completed after calculating mass error (**Table S4-S8**) of three biological replicates (N=3) and signals were only considered significant if altered regulation was observed in two or more replicates.

### PQS complementation

The complementation strains *P. aeruginosa* PA14 Δ*pqsA-C::pqsA-C*, Δ*pqsH::pqsH*, and Δ*pqsL::pqsL* were constructed as described in Jo *et al.* [57]. Primers LD1 & LD4, LD168 & LD171, and LD9 & LD12 were used to amplify the *pqsA-C, pqsH* and *pqsL* genes, respectively (**Table S10**). Correct constructs were confirmed by PCR and sequencing and complemented into the original deletion site, following the same procedure as for deletion.

### Bacterial Extraction for PCA and PCH Fold-change

Bacterial growth for quantification proceeded exactly as described in “Imaging mass spectrometry experiments”. Each strains (PA14 WT, Δ*pqsA-C*, Δ*pqsH*, Δ*phz*, Δ*pqsA-C::pqsA-C* and Δ*pqsH::pqsH*) was plated in duplicate. After 48 hours, a spatula was used to separate the entire colony and surrounding agar from petri dish. Samples were transferred to a glass vial. Each vial was filled with 2-3 mL of DI water and homogenized using a spatula. Vials were either flash frozen or stored at −80□ prior to lyophilization. Upon dryness, 3mL of MeOH was added to each vial and samples were sonicated for 30 minutes to extract organics. Total weights of each sample dry biomass were used to achieve a 10mg/mL solution of each extract. 10mg/mL solutions were centrifuged at 10,000 rpm for 2 minutes before transferring to HPLC vials for analysis. (Repeated for a total of two biological replicates).

### PCA and PCH Fold-change analysis

PCA standard was purchased from ChemScience (>97%). An HPLC method previously described by *Adler et al.* was used on an Agilent 1260 Infinity to isolate and identify PCA through retention time matching with standard (32.5 min at 250 nm; **Figure S7**) with a Phenomenex C18 analytical column (150 × 4.6 mm; 5 μm) and a flow rate of 0.5 mL/min [27]. Area under the curve (AUC) was used to quantify fold-change between PA14 WT and other strains/conditions. For PCH fold-change quantification, a gradient of 10%-85% ACN (0.1% TFA) and H20 (0.1% TFA) over 25 minutes with the two PCH stereoisomers (pyochelin I and II) eluting at 16.0 & 16.2 minutes respectively at 210 nm. Area reported for fold change analysis was achieved by combining the AUC of the peaks for pyochelin I and II. Fold-change analysis was determined averaging the areas of EICs for two independent biological cultures (N=2), with three technical replicates (n=3), and compared to PA14 WT with no TLCA (control).

## Results & Discussion

Phenazines constitute one of the most notable families of specialized metabolites produced by *P. aeruginosa.* Phenazines are redox-active compounds that have been implicated in balancing redox homeostasis in the hypoxic regions of biofilms, thereby regulating biofilm morphology [28–32]. A phenazine-null mutant (Δ*phz*) over produces extracellular matrix which causes Δ*phz* biofilms to have a characteristic hyper wrinkled morphology that allows for increased access to oxygen due to a higher surface area-to-volume ratio [32–35]. To characterize the effect of TLCA on *P. aeruginosa* biofilms, we grew PA14 colonies in a colony morphology assay on solid agar supplemented with Congo Red, Coomassie Blue, and TLCA (0, 100, 250, and 1000 μM) for five days (**Figure S1**). Wild type PA14 initially forms a smooth colony but initiates wrinkle structure development after 70 hours (**Figure 1**). In contrast, a Δ*phz* mutant showed enhanced production of biofilm matrix, leading to an earlier onset of wrinkling between 24 and 48 hours. Addition of TLCA to the medium promoted a hyper-wrinkled morphology after 94 hours. The colony continued to wrinkle over time, resembling the Δ*phz* mutant (118 hours; **Figure 1**). This morphology phenotype was also observed at lower concentrations, but it was most dramatic at 1 mM TLCA, which is a physiologically relevant concentration [36] (**Figure S1**). Based on this assay, it appears that TLCA induces matrix production, possibly by downregulating phenazines.

**Figure 1:**
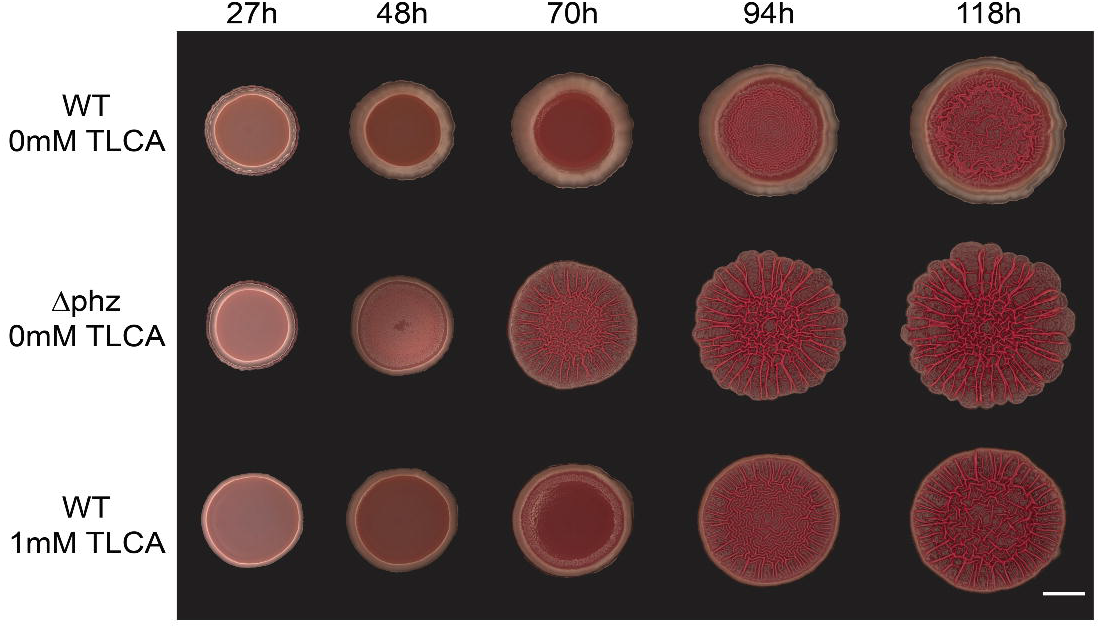
The effect of TLCA on colony biofilm formation in *P. aeruginosa* PA14. After five days of growth, colonies that were exposed to TLCA showed a similar hyper-wrinkled biofilm structure to that of the untreated Δ*phz* mutant.

The putative decrease in phenazine production as indicated by the Δ*phz*-like colony morphology in the presence of TLCA was queried alongside other changes in specialized metabolism using imaging mass spectrometry (IMS). We employed matrix-assisted laser desorption/ionization time-of-flight IMS (MALDI-TOF IMS), because it provides a robust, untargeted analysis of the specialized metabolites produced by *P. aeruginosa in situ* [37,38]. *P. aeruginosa* colonies were grown on thin agar (2-3 mm) embedded with the vehicle or 250 μM of TLCA for 48 hours. The colonies and their respective agar controls were then prepared for IMS analysis. Twelve known specialized metabolites were identified and visualized from *P. aeruginosa* colonies (**Figure 2**). Orthogonal analytical techniques were used to confirm identities of all twelve metabolites (**Figure S2**). Following a combination of manual and statistical analyses using SCiLS lab, eight of the twelve specialized metabolites were observed to have significant altered regulation in the presence of TLCA (p<0.05; **Table S1**). The identified specialized metabolites represent four broad classes of molecular families including the phenazines, quinolones, rhamnolipids, and siderophores. We found that the phenazines pyocyanin (PYO) and phenazine-1-carboxamide (PCN) are significantly downregulated when TLCA is present, supporting our hypothesis that TLCA exposure causes hyper-wrinkled colonies by downregulating phenazine production. We did not observe a statistically significant change in phenazine-1-carboxylic acid (PCA) production in the presence of TLCA. Since methylated phenazines like PYO, but not PCN and PCA were shown to inhibit colony wrinkling [39] our data is consistent with TLCA affecting colony morphology by modulating phenazine production.

**Figure 2:**
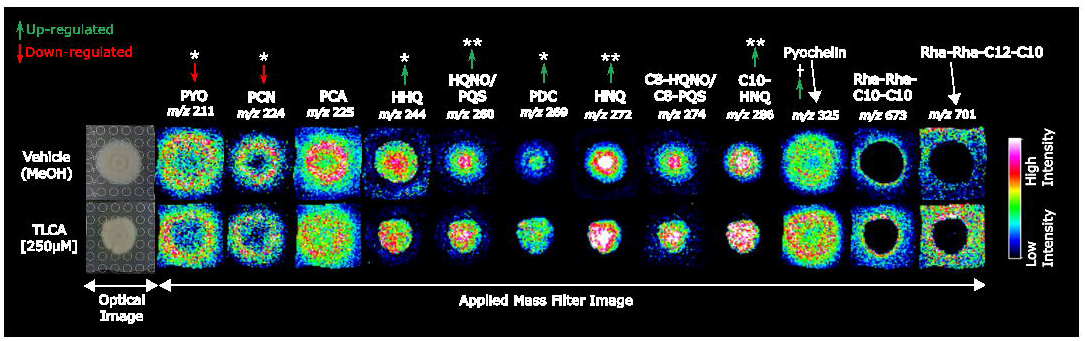
MALDI-TOF IMS analysis of *P. aeruginosa* after exposure to TLCA. Twelve characterized specialized metabolites produced by *P. aeruginosa* were identified and visualized. Signal intensity is displayed as a heat map and shows that exposure to TLCA altered regulation of highlighted specialized metabolites compared to control. * denotes the signal is significantly up- or down-regulated in two biological replicates within the colony and □ denotes significance in the surrounding agar (p<0.05). Two asterisks (**) denote the signal was significant over all three biological replicates.

*P. aeruginosa* is reported to produce up to 50 quinolones which are specialized metabolites that play specific roles in signaling and/or virulence. For example, both 2-heptyl-4-quinolone (HHQ) and *Pseudomonas* quinolone signal (PQS) are specifically known for their signaling properties but have also demonstrated antifungal bioactivity [40–46]. The N-oxide quinolone, 4-hydroxy-2-heptylquinoline-N-oxide (HQNO), has recently been shown to have antimicrobial activity towards Gram-positive bacteria and contributes to *P. aeruginosa*’s virulence [40–45]. When exposed to TLCA, we observed a significant upregulation of HHQ, PQS, and 4-hydroxy-2-nonylquinoline (HNQ) within the colony. PQS is a well-characterized signaling molecule in *P. aeruginosa* quorum sensing cascade [47]. Though IMS is a valuable tool for identifying and visualizing the chemical composition of a sample, it cannot differentiate between constitutional isomers like HQNO and PQS (*m/z* 260; **Figure 2**). In order to differentiate these two metabolites, we used a combination of tandem mass spectrometry and knockout mutants to demonstrate that PQS was represented by the signal that is retained in the colony (center, upregulated) and HQNO corresponds to the signal that was excreted into the agar (outer signal, downregulated) (**Figure S3**). HHQ and PQS are well established signaling molecules that are required for phenazine production in *Pseudomonas aeruginosa*, However, since we observed decreased phenazine levels despite an increase in quinolone production [48] our results suggest that TLCA is attenuating phenazine production in a quinolone-independent manner.

Based on the IMS analyses, treatment of TLCA seems to induce the production of biosurfactants. These compounds, such as rhamnolipids, are amphipathic small molecules that *P. aeruginosa* produces to increase surface adhesion and motility [49]. Rha-Rha-C10-C10 and Rha-Rha-C12-C10/C10-C12 production were produced at elevated levels after treatment with TLCA (**Figure 2**). However, while these results were not statistically significant, the trend is worth noting because increases in rhamnolipid production would agree with the previously reported bioactivity of TLCA. In colony biofilms, TLCA markedly increases matrix production and leads to increased spreading and wrinkling. This wrinkly spreader phenotype is reminiscent of the phenazine-null mutant as seen in **Figure 1**.

In addition to quinolones and phenazines, we detected changes in the levels of the siderophore pyochelin. Siderophores are iron chelators that allow bacteria to acquire iron from the surrounding environment [50]. Siderophores have been known to sequester iron from host proteins and simultaneously act as signals for biofilm development [12]. The IMS results show a significant upregulation of pyochelin in the presence of TLCA (**Figure 2**). Pyochelin is produced by the biosynthetic pathway *pchA-I* which is activated by the presence of both iron and the ferric uptake regulator (Fur) [12]. Additionally, previous work has shown that an increase in iron-bound PQS can indirectly increase siderophore production by activating the siderophore gene clusters *pvd* and *pch*. Our IMS experiments on wild-type PA14 (**Figure 2**) confirm this relationship due to the increase of both PQS and pyochelin in the presence of TLCA. Siderophores have antimicrobial activity and contribute to virulence [51,52]. Hence, the TLCA-dependent upregulation of pyochelin raised the question of whether TLCA-exposed *P. aeruginosa* become hypervirulent.

Many bacteria, like the ESKAPE pathogens, exhibit distinct lifestyle states depending on surrounding environmental factors. Chua *et al.* recently described characteristics of the dispersed cell state using sodium nitroprusside (SNP) as a biofilm-dispersing agent [22]. They found that dispersed cells are characterized by altered physiology, increased virulence against macrophages and *C. elegans*, and extreme sensitivity to iron starvation [22]. Having observed an increase in pyochelin production when exposing *P. aeruginosa* to TLCA (**Figure 2**), we sought to determine if TLCA treated cells were hypervirulent using a *Galleria mellonella* (greater wax moth) larvae virulence assay.

*G. mellonella* larvae have been shown to be an ideal model for studying microbial pathogenesis of several ESKAPE pathogens since they are easily infected, inexpensive, and produce a similar immune response as vertebrates and mammals [53–57]. Larvae were injected with a *P. aeruginosa* or vehicle load, and exposed to SNP, TLCA, or no dispersing agent two hours post infection. In the infected TLCA-exposed larvae, a 50% decrease in survival rate was observed 17 hours post infection compared to the infected larvae that did not receive a dispersing agent (**Figure 3**). In the *G. mellonella* model, the potency of treatment is gauged by the survival curve, therefore a shift in this curve will indicate increased or decreased virulence of *P. aeruginosa* as compared to the 50% survival rate of non-treated infected larvae [58].

**Figure 3:**
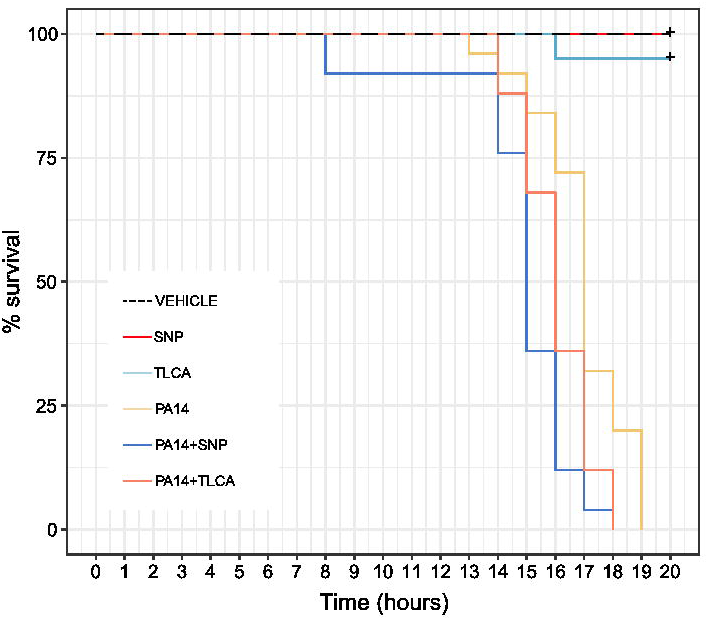
Virulence assay with *Galleria mellonella* (greater wax moth). Using *G. mellonella* as an infection model revealed that regardless of biofilm dispersing agent, dispersed biofilm cells show a significant increase in virulence compared to controls (p<0.05).

Addition of SNP or TLCA lead to a significant decrease in survival with 50% of the larvae succumbing to infection 1-2 hours earlier than the PA14 control. Exposure to either of the biofilm inhibitors, SNP or TLCA, did not affect the survival rate of uninfected larvae. TLCA-dispersed cells are also hypervirulent and have an analogous trend to SNP-dispersed cells. This confirms that TLCA-dispersed cells are significantly more virulent than planktonic or biofilm cells as previously observed with SNP-dispersed cells (**Table S3**; p<0.05).

Having confirmed that TLCA increases virulence in *P. aeruginosa,* we were interested in testing whether *P. aeruginosa* TLCA-treated cells would also be sensitive to iron stress and whether this may be related to the increase in pyochelin production. In SNP-dispersed cells, pyoverdine the other main siderophore produced by *P. aeruginosa,* was markedly decreased and these dispersed cells showed a sensitivity to iron starvation [22]. However, we were unable to detect pyoverdine in either the wild type or TLCA-treated *P. aeruginosa* via mass spectrometry, but were able to detect pyochelin which Chua *et al.* did not measure (Figure 2). Therefore, we recapitulated the iron starvation assay performed by Chua *et al.* to determine if TLCA exposure induces sensitivity to iron starvation as shown for SNP-dispersed cells.

Iron starvation was induced by exposing PA14 cells to the iron chelator 2’2-bipyridine (DIPY; **Figure 4**). This assay was performed with TLCA-dispersed biofilm cells, which were generated from pellicles (biofilms grown at the air-liquid interface) (**Figure 4A & B**), and planktonic cells, where exposure to TLCA acted to inhibit biofilm formation (**Figure 4C & D**). Proliferation of the cells was monitored by measuring optical density at 600 nm of each condition over 16 hours. TLCA exposure alone does not alter proliferation compared to the untreated control (**Figure 4B**). While exposure to DIPY, which induces iron starvation, significantly decreases proliferation (p<0.05) of biofilm-derived cells during the log phase of growth (120-540 mins) (**Figure 4B**). However, there is no significant difference between biofilm cells and TLCA-dispersed biofilm cells when exposed to iron starvation. We observe the same trends when inhibiting biofilm formation of planktonic cells with TLCA (**Figure 4D**). Therefore, unlike for SNP, TLCA-dispersed cells are not sensitive to iron starvation.

**Figure 4:**
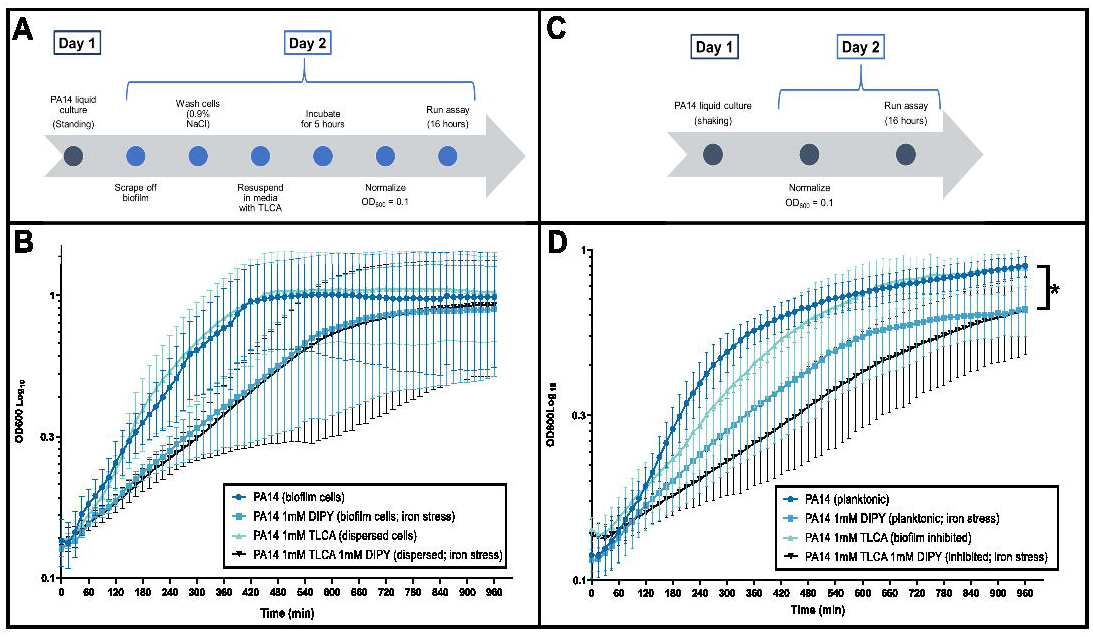
PA14 Iron starvation assay. (A & B) shows that when biofilm dispersed *P. aeruginosa* cells are exposed to TLCA there are no changes in proliferation. When exposed to iron starvation, we observe no significant difference between biofilm and dispersed biofilm cells. Confirming TLCA dispersed cells are not more sensitive to iron starvation than biofilm cells. (C & D) We observe the same relationship with planktonic cells and planktonic cells inhibited from entering the biofilm state. However, planktonic cells do show a sensitivity to iron starvation. * denotes p<0.05.

The discrepancy in iron starvation sensitivity might be attributed to their different effects on pyoverdine production. Though we were unable to detect pyoverdine in our IMS analysis, iron starvation might be prevented by the increased presence of pyochelin in colony biofilms that were exposed to TLCA (Figure 2). Despite their differing effects on iron starvation sensitivity, both SNP and TLCA have previously been shown to readily disperse biofilms, likely through different mechanisms of action [18,22]. SNP, a nitric oxide donor, has been shown to disperse mature biofilms by producing nitrosative stress inside of the biofilm structure [21]. Since TLCA cannot act as a nitric oxide donor and it does not cause cell lysis, TLCA must act via an alternative mechanism to disperse biofilms [18]. This, along with results, supports our hypothesis that TLCA exposure induces alterations to *P. aeruginosa’s* specialized metabolism.

PQS is a major quorum sensing signal and iron-bound PQS upregulates siderophore production [59]. The increase in pyochelin production *in situ* and the enhanced virulence *in vivo* lead us to investigate the contribution of the *pqs* gene cluster to the TLCA-mediated effect. We tested four mutants with deletions in quinolone and phenazine biosynthetic genes: Δ*pqsA-C*, Δ*pqsH*, Δ*pqsL*, and Δ*phz*. Δ*pqsA-C* does not produce any quinolones, while Δ*pqsH* cannot produce PQS and Δ*pqsL* cannot produce N-oxide quinolones such as HQNO [48,60,61]. The phenazine-null mutant, Δ*phz,* is a double-deletion of the two redundant core phenazine biosynthetic gene clusters *phzA1-G1* and *phzA2-G2* [48,60,61]. Using IMS, we investigated if any of the four mutants would recapitulate the TLCA-dependent increase in pyochelin production that we observed for the WT. No variation in pyochelin production for Δ*pqsA-C* and Δ*pqsL* mutants were found, while Δ*pqsH* mutant produced significantly more pyochelin in response to TLCA, mimicking the trend observed in the WT (p<0.05; **Figure S6**). This result puts into question our earlier assumption that PQS and pyochelin production are positively correlated. We also observed a significant increase in pyochelin production in the Δ*phz* mutant (**Figure S6**). Previous work has shown that increasing PCA concentrations allows PA14 siderophore-null mutants to still develop biofilms and sequester iron [62]. This may be due to phenazine’s ability to mediate the reduction of Fe(III) to the bioavailable Fe(II) [62]. *pqsH* and *phz* gene clusters are necessary for phenazine production, hence a decrease or lack of phenazine production due to TLCA exposure might lead to the observed increase in pyochelin production (**Figures 2 & S6**).

Since our IMS results were inconclusive regarding the effect of TLCA on PCA production in colony biofilms (**Figure 2**), we subjected bacterial colony extracts from wild-type PA14, Δ*pqsH*, and Δ*phz* to an HPLC analysis to measure fold change across biological (N=3) and technical replicates (n=3) (**Figure 5A**). When considering fold changes greater than +/− 1, PCA production was not altered by TLCA exposure in the wild type and Δ*pqsH* strains (not produced by Δ*phz* mutant). However, an increase in pyochelin production was observed in the TLCA-exposed Δ*phz* mutant (+2.20) (**Figure 5B**). When the Δ*pqsH* mutant is complemented (Δ*pqsH::pqsH*) there is increased production of pyochelin compared to wild-type whether the colony is TLCA-treated or not (+2.43 and +1.81, respectively) (**Table S9**). There was no notable difference in production of pyochelin in the complementation strain whether it was TLCA-treated or not, which recapitulates the wild-type data. Though the fold change of PCA and pyochelin was comparable in both wild-type PA14 and Δ*pqsH,* the change in the production of these two metabolites in the phenazine-null mutant, Δ*phz,* suggests that a lack of phenazine production is correlated to the observed increase in pyochelin production from cells exposed to TLCA.

**Figure 5:**
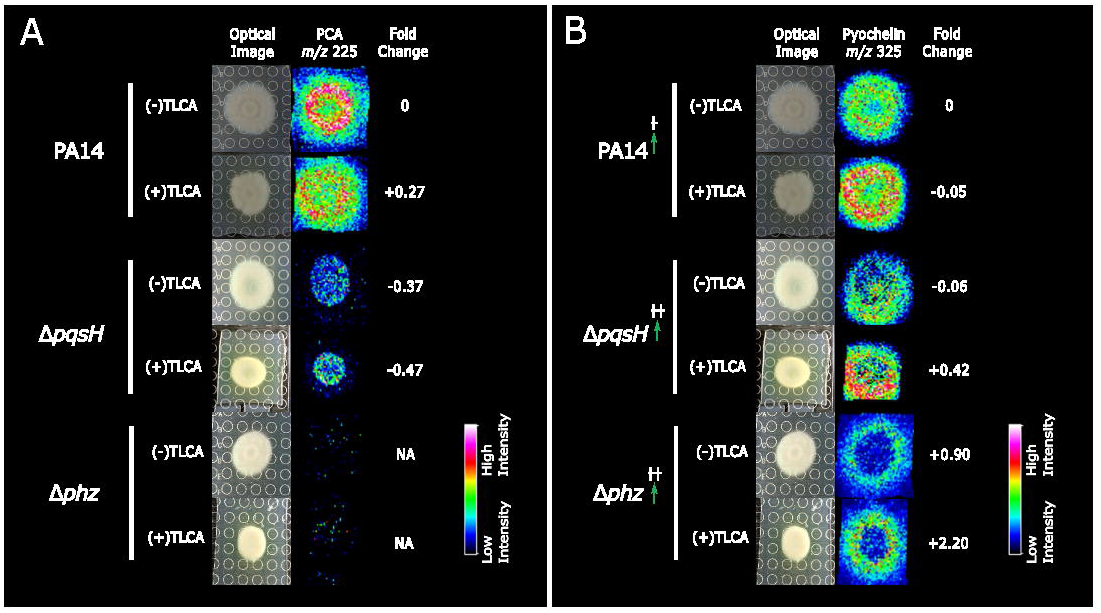
IMS and fold change analysis of wild type PA14, *pqsH, and phz* mutants. (A) IMS analysis revealed TLCA exposure induced no change in PCA production however (B) there was an increase in pyochelin production in the Δ*phz* mutant, resembling the trend observed in wild-type experiments. □□ denotes the observed regulation was statistically significant over three biological replicates in IMS experiments (p<0.05).

## Conclusion

In this study we demonstrate that the endogenous enteric biofilm inhibitor, TLCA, can alter colony morphology, specialized metabolism, and virulence of *P. aeruginosa.* Our biological and chemical studies confirm what is already known about TLCA’s bioactivity and offers insight into the chemical communication occurring between the cells upon treatment with a known biofilm inhibitor. TLCA-dispersed biofilm cells are not sensitive to iron starvation, as previously reported for the biofilm-dispersing agent SNP, however dispersion via either agent induces a significant increase in virulence *in vivo,* implying that the mechanism of action of the two biofilm inhibitors is different. Our IMS analysis of mutant strains revealed that when exposed to TLCA, a lack of PQS (Δ*pqsH*) or phenazine production (Δ*phz*) lead to an increase in pyochelin production, matching the results observed in our colony morphology and IMS WT experiments. Therefore, PQS or iron-bound PQS is not responsible for activating the *pchA-I* gene cluster (**Figure 6A**). However, IMS analysis highlighted that a significant increase in pyochelin production was observed in the Δ*phz* mutant confirming that the *phz* gene cluster plays a role in increased pyochelin production when TLCA is present (**Figure 6B**). Though these results are not conclusive regarding the mechanism of TLCA dispersal since biofilm inhibition and dispersal cannot be tested using colony biofilms as a model system, it does support the hypothesis that TLCA is acting as an environmental cue to induce *P. aeruginosa* to alter its metabolic signaling throughout the bacterial community, thus disrupting the biofilm life cycle.

**Figure 6:**
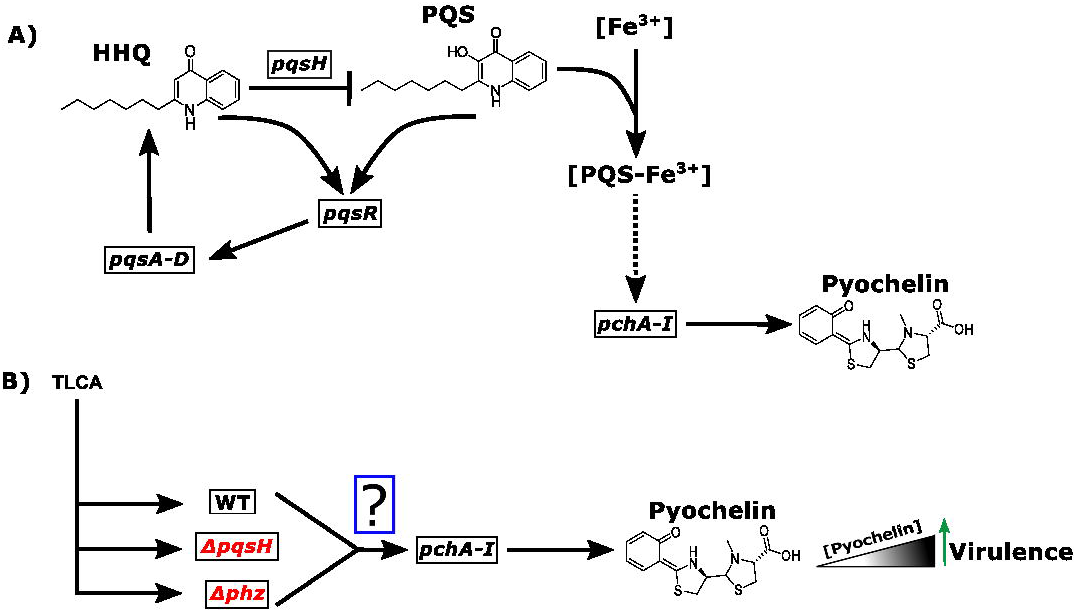
*pqs* metabolic pathway under TLCA exposure. (A) Represents the canonical pathway for pyochelin production with the role iron-bound PQS plays in indirectly (dashed-line) activating pyochelin production. (B) However, TLCA treatment of the WT, *pqsH,* and *phz* mutants leads to an increase in pyochelin production and subsequently virulence supporting that these genes play a role in the observed increase in pyochelin production from TLCA exposure.

More work is needed to determine the mechanism of action of TLCA, but our work shows that through TLCA exposure *P. aeruginosa* bacterial communities increase the production of virulence factors such as pyochelin and consequently develop hypervirulence *in vivo.* Others have documented similar phenomena by reporting an increase in virulence *in vivo* as a consequence of exposure to a biofilm inhibitor in organisms such as, Group A *Streptococcus, Vibrio cholerae,* and *Candida albicans* [63–67]. Taken into consideration previous literature, our work promotes that future investigations must consider how biofilm inhibitors alter the chemical environment within a bacterial biofilm and that biofilm inhibition as a treatment strategy should be closely monitored for variations in specialized metabolism leading to undesired side effects such as increased virulence.

## Acknowledgements

We thank Drs. Atul Jain and Terry Moore for assisting with the recrystallization of the MALDI matrices. Funding was provided by Grant K12HD055892 from the National Institute of Child Health and Human Development (NICHD) and the National Institutes of Health Office of Research on Women’s Health (ORWH) (L.M.S.); University of Illinois at Chicago Startup Funds (L.M.S.); American Society for Pharmacognosy research startup grant (L.M.S.); NIH/NIAID grant R01AI103369 and an NSF CAREER award (L.E.P.D). ARC was supported in part by the National Science Foundation Illinois Louis Stokes Alliance for Minority Participation Bridge to the Doctorate Fellowship (grant number 1500368) and a UIC Abraham Lincoln retention fellowship.

