## Supplemental Information for "Biofilm inhibitor taurolithocholic acid alters colony morphology, specialized metabolism, and virulence of *Pseudomonas aeruginosa*"

| 1. General experimental methods……………………………………………………………. | S3 |
| --- | --- |
| 2. Figure S1: Colony morphology studies with 250 µM and 1 mM TLCA………………... | S5 |
| 3. Figure S2: Molecular networking of *P. aeruginosa* specialized metabolism………….. | S6 |
| 4. Figures S3: Replicate One of IMS images of PA14 characterized specialized metabolites…………………………………………………………………………………….... | S9 |
| 5. Figures S4: Replicate Two of IMS images of PA14 characterized specialized metabolites…………………………………………………………………………………….... | S10 |
| 6. Figures S5: Replicate Three of IMS images of PA14 characterized specialized metabolites…………………………………………………………………………………….... | S11 |
| 7.Figure S6: PA14 Mutant IMS analysis of PCH Production ……………………………... | S12 |
| 8. Figure S7: Retention time matching and UV-vis profile of PCA………………………... | S13 |
| 8. Figure S8: GNPS search results of PCH for metabolite identification……………….... | S14 |
| 9. Figure S9: UV-vis profile of isolated PCH for metabolite fold-change analysis………. | S15 |
| 11. Table S1: Statistically significant signals from PA14 IMS experiments………………. | S16 |
| 12. Table S2: Mass error for PA14 IMS experiments………………………………………. | S17 |
| 13. Table S3: Statiscal analysis of wax moth virulence assay…………………………….. | S18 |
| 15. Tables S4: Statistically significance of pyochelin signals from mutant IMS experiments…………………………………………………………………………………….. | S19 |
| 16. Tables S5: Mass error from Δ*pqsA-C* IMS experiments………………………………. | S20 |
| 17. Tables S6: Mass error from Δ*pqsH* IMS experiments…………………………………. | S21 |
| 18. Tables S7: Mass error from Δ*pqsL* IMS experiments………………………………….. | S22 |
| 19. Tables S8: Mass error from Δ*phz* IMS experiments…………………………………… | S23 |
| 20. Table S9: Fold Change analysis of PCA and PCH from TLCA treatment….............. | S24 |
| 21. Table S10: List of strains, plasmids, and primers used………………………………... | S25 |
| 22. Data Availability………………………………………………………………………….... | S28 |

*Large scale cultivation and crude extract preparation.* An overnight 5 mL liquid culture was transferred to 50 mL of LB liquid media in a 250 mL Erlenmeyer flask, covered with a KenAG milk filter and autoclave paper, and left to shake overnight at 30 ℃. The following day, the 50 mL liquid culture was transferred to 1 L of LB media in a 2.8 L Fernbach flask with milk filter and autoclave paper and left to shake for four days at 30℃. Two growth conditions were employed, 1) a qualitative control where the liquid culture was left to shake for four days and then extracted on day five and 2) exposure to TLCA where an aliquot of TLCA was added to the 1L liquid culture for a final concentration of 250 µM of TLCA after overnight growth at 30 ℃ and left to shake for four days before extraction (Extraction on day 5 of the liquid culture with 4 days of TLCA exposure). To generate crude extracts 20 G of Amberlite XAD16N resin (Sigma) was added to the culture and allowed to shake at 225 rpm at 30 ℃ for one hour. The cells and resin were filtered with filter paper (Grade 2/ 8 µM poresize & Grade 934-AH/ 1.5µM pore size, respectively) and filter paper and resin was transferred to a Erlenmeyer flask. Resin was back extracted using 250 mL of methanol and flask was placed back on shaker at 30 ℃ for an hour. The resin and methanol solution was then filtered and the organic extract was dried *in vacuo* to yield a crude organic extract. This crude extraction was subject to solid phase extraction via a Supleco Discovery DSC-18, 5 g, 20 mL tube to separate the crude extract into seven rarified extracts based on polarity (A: 10% MeOH, B: 20% MeOH, C: 40% MeOH, D: 60% MeOH, E: 80% MeOH, F: 100% MeOH, G: 100% EtoAc). The seven collected fractions were then dried down *in vacuo* and stored at -20 ℃.

*LC/MS-MS preparation and experimentation.* Each of the seven fractions (A-G) were subjected to LC-MS/MS analysis. Each sample was dissolved in LC-MS grade MeOH to a final

concentration of 400 µg/mL. All samples were subjected to the same LC method (23 minute gradient from 20% MeOH to 95% MeOH with a 10 minute wash and 10 minute equilibration). A Phenomenex Kinetex 5 µm C18 100 **Å** 150 x 4.6mm was used for separation on an Agilent LC-1200 dual-pump HPLC instrument. Separated samples were then sent to a Thermo Finnigan LCQ Advantage Max LC-MS/MS for fragmentation with an ESI source via data dependent fragmentation. Settings for fragmentation data: temp 250 ℃, voltage 35 eV, isolation window 2 Da, collision energy 35%, three MS^2^ instances for every MS^1^.

*GNPS & Molecular Networking*. For molecular networking analysis raw tandem MS data was uploaded to GNPS (Global Natural Product Social Molecular Networking). GNPS is an open access knowledge database for the comparison of raw MS/MS data to a library of MS/MS data of known natural products [[1]](https://paperpile.com/c/xWBxHG/YJaj). For this study, all libraries in GNPS were used for comparison and selected for this workflow under “Spectral Library”. In “basic options” precursor ion mass tolerance was set to 0.5 Da and fragment ion mass tolerance was set to 0.3 Da. Under “advanced network options” the minimum matched fragment ions was set to 4. For “advanced library search options” the library search min matched peaks was also changed to 4. Under “advanced filtering options” the filter precursor window was set to “Don’t Filter” whereas filter library option was left as defaulted showing “Filter Library”. All other options are left at the default settings. Using cytoscape 2.8.3 the data was analyzed with FM3 layout [[2]](https://paperpile.com/c/xWBxHG/tzkp).

*Pyochelin (PCH) Identification.* For PCH fold-change quantification, a gradient of 10%-85% ACN (0.1% TFA) and H_2_0 (0.1% TFA) over 25 minutes where separated samples were then sent to a Thermo Finnigan LCQ Advantage Max LC-MS/MS for fragmentation with an ESI source via data dependent fragmentation. Settings for fragmentation data: temp 250 ℃, voltage 35 eV, isolation window 2 Da, collision energy 35%, three MS2 instances for every MS1. After identifying the UV peak and retention time which correlated to PCH (*m/z* 325 [M+H]^+^, RT: 18.9) in our PA14 WT extract (N=2). Fragmentation data was uploaded to GNPS and confirmed as a positive hit for the metabolite pyochelin (**Figure S8**).

**Figure S1**: Colony morphology study of *P. aeruginosa* exposed to two TLCA concentrations (250 µM and 1 mM) over 120 hours. An increase in wrinkling phenotype is observed compared to control with altered morphology correlating to an increase in TLCA concentration.

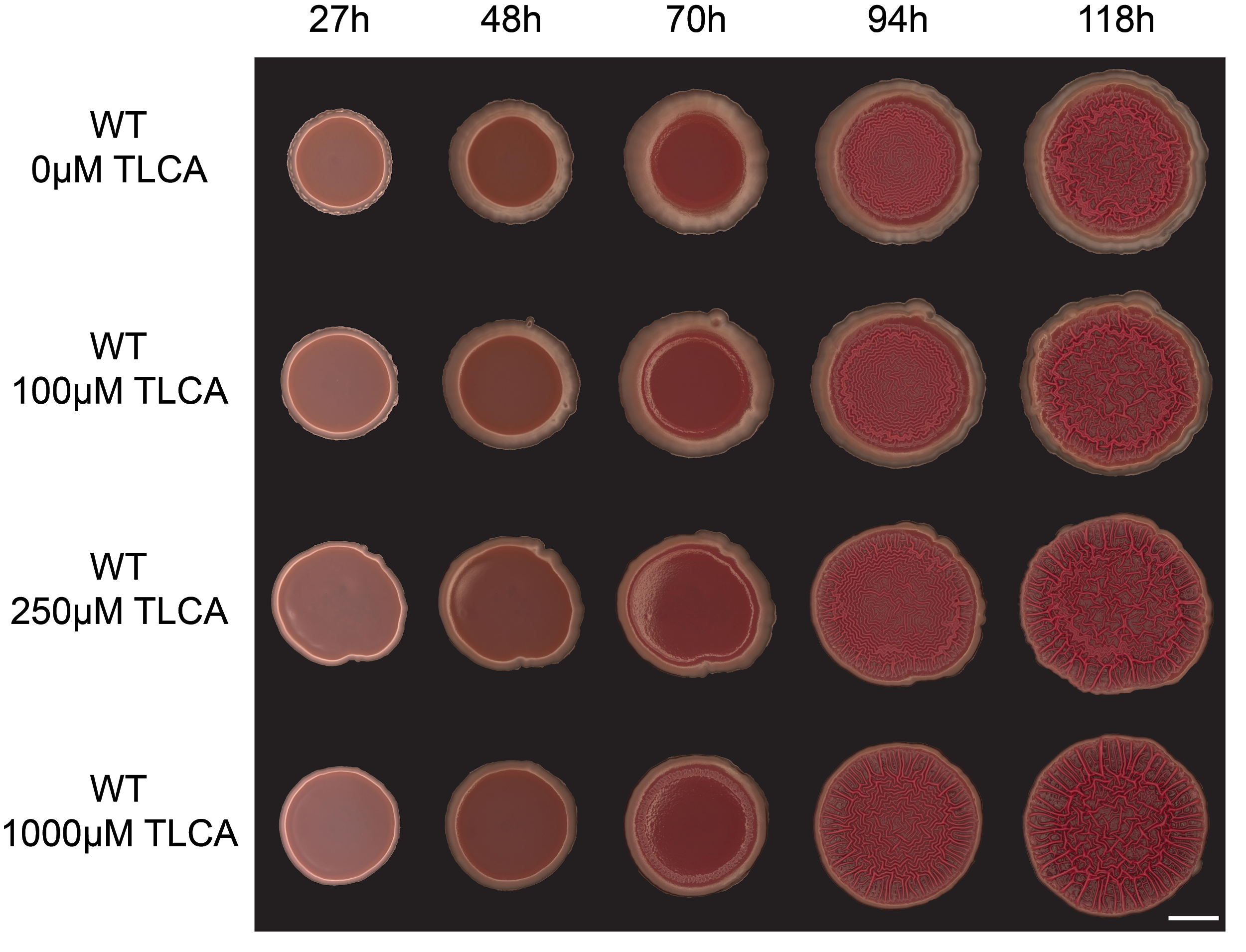

**Figure S2**: Complete molecular networks of extracted metabolites from *P. aeruginosa* exposed to TLCA. All background noise such as nodes corresponding to only TLCA or media controls were removed prior to analysis. The network on the left is used to highlight the smaller metabolites of group A (200-230 Da) due to the smaller size of the metabolites and therefore different settings required for GNPS analysis. Both networks represents a total of 1,029 with 181 of the nodes representing metabolites only found in the TLCA condition. The network on the right represents three technical replicates of LC-MS/MS data. Molecular network clusters of (A) phenazine family, (B) quinolone family, (C) rhamnolipids, and (D) the siderophore pyochelin with respective cosine values.

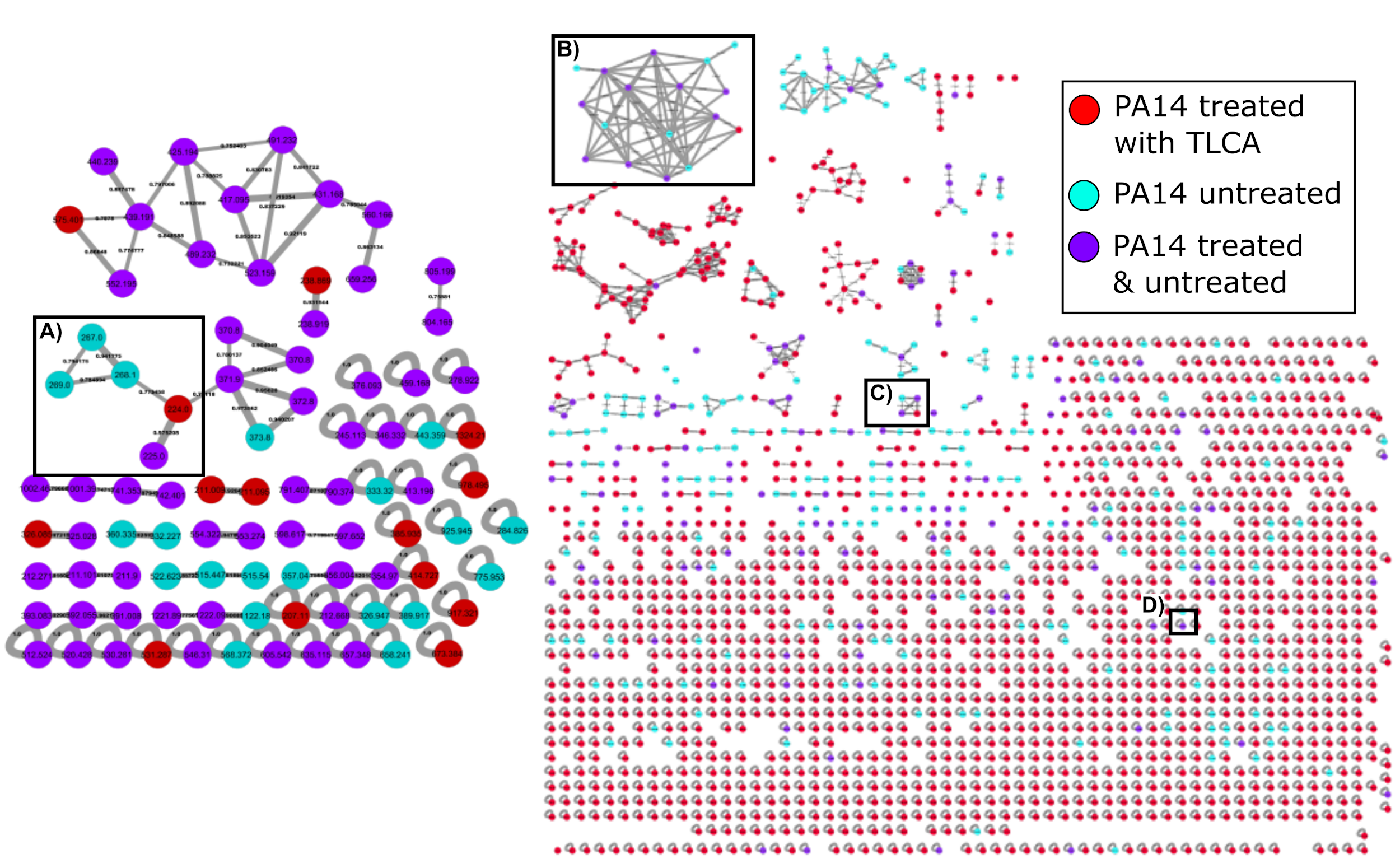

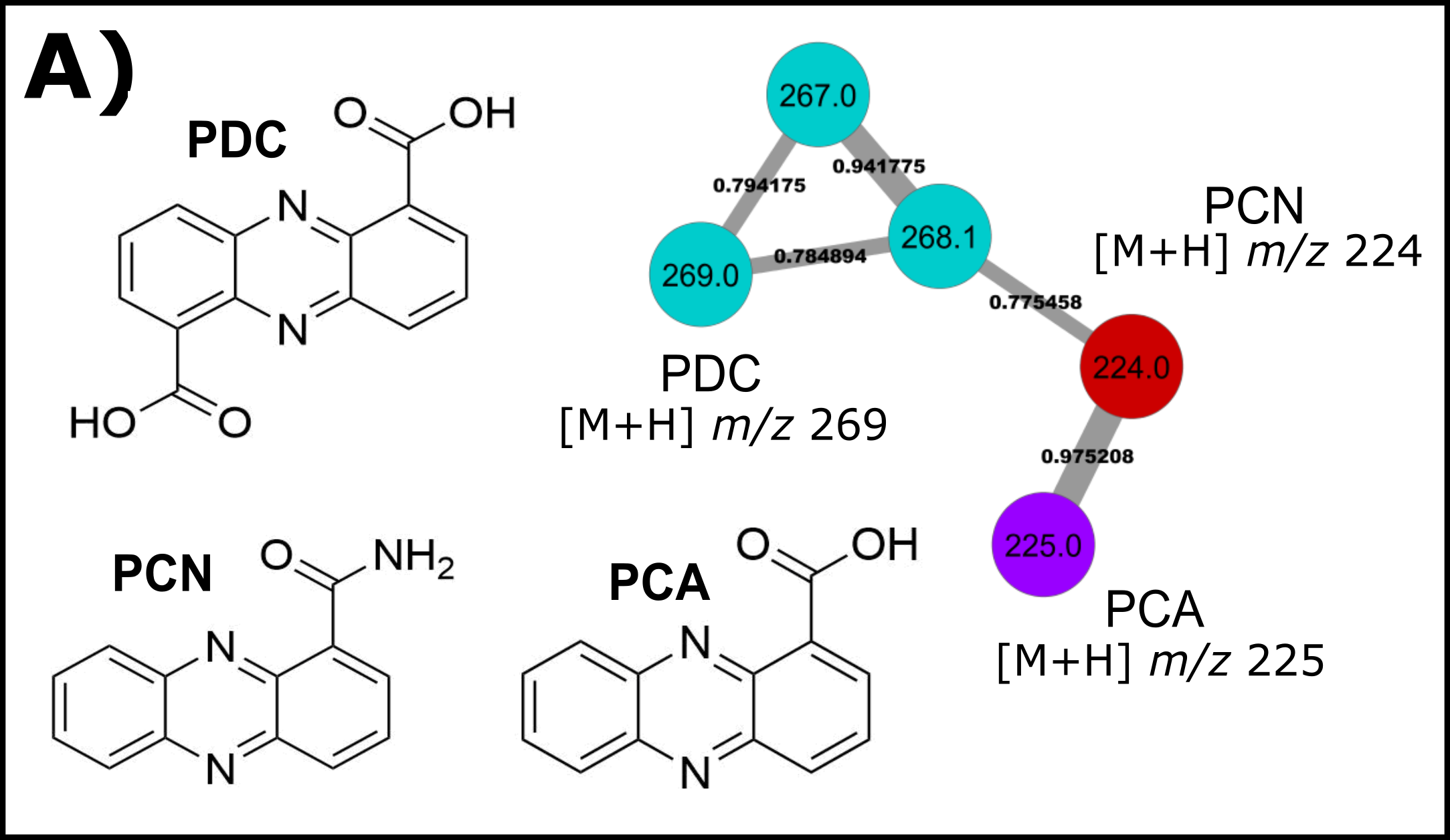

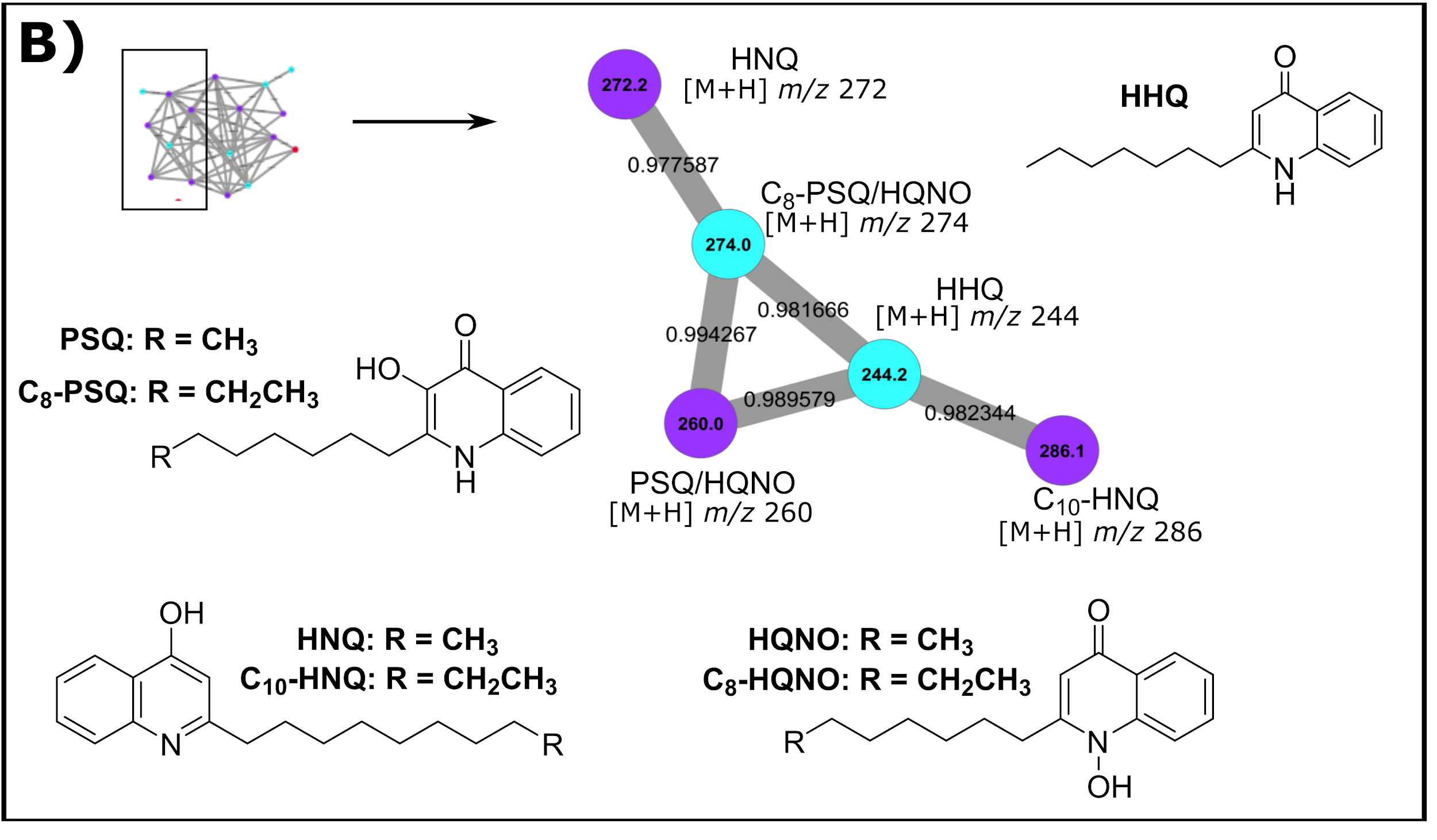

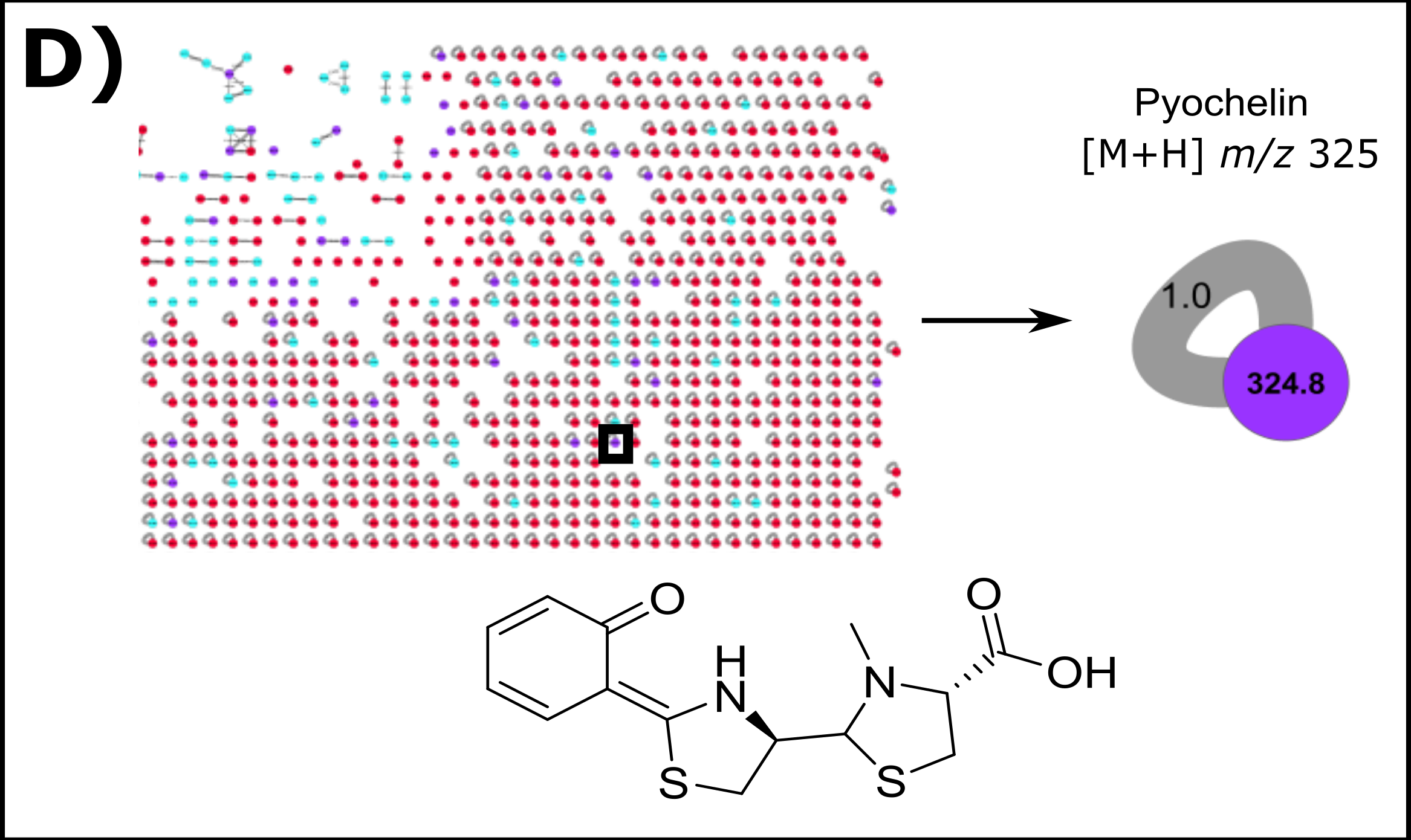

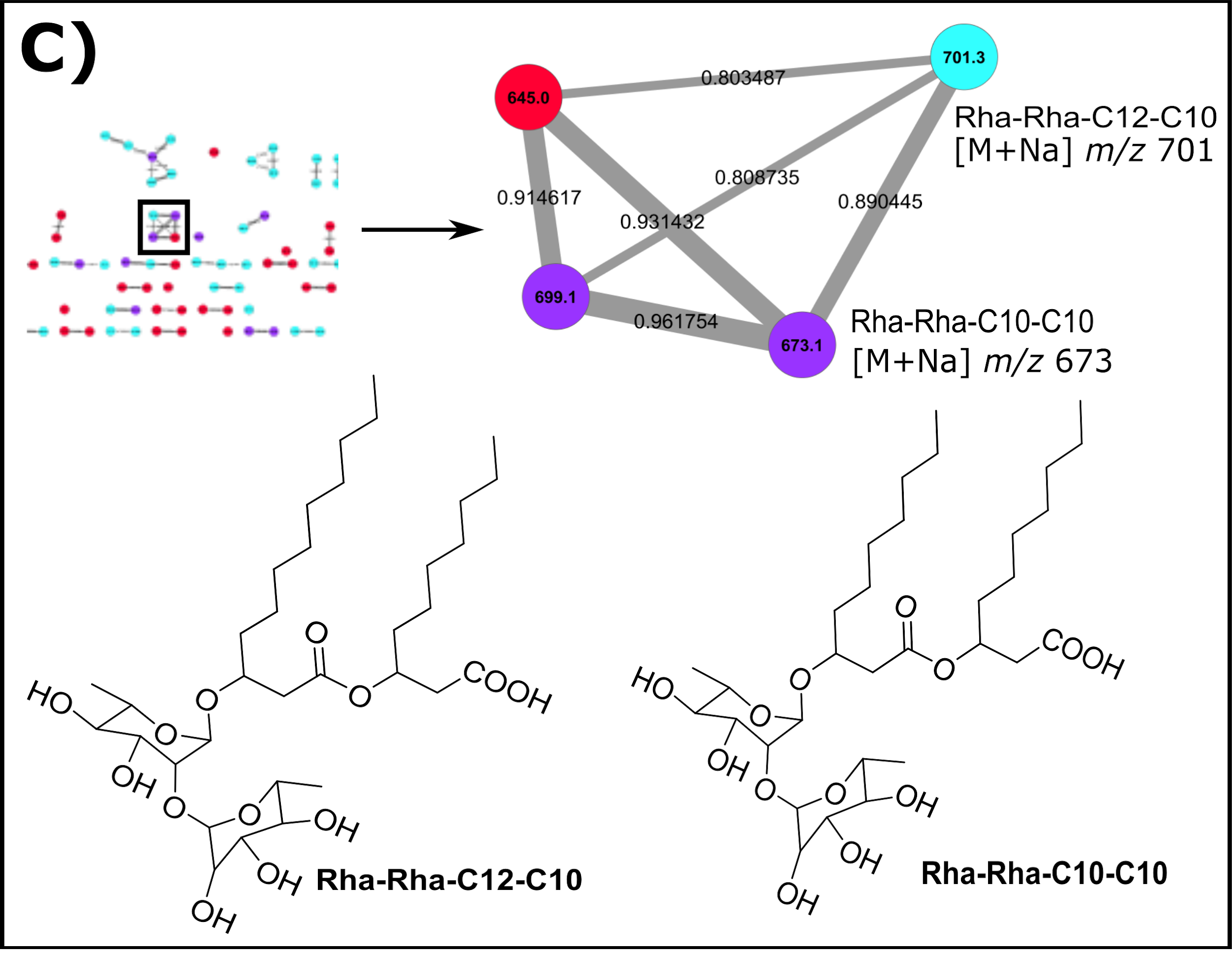

**Figure S3**: Replicate one IMS analysis of the twelve characterized metabolites seen in Figure 2. Statistical analysis reveals where a signal has altered regulation within the colony (*) or in the surrounding agar (ᵻ) (p<0.05).

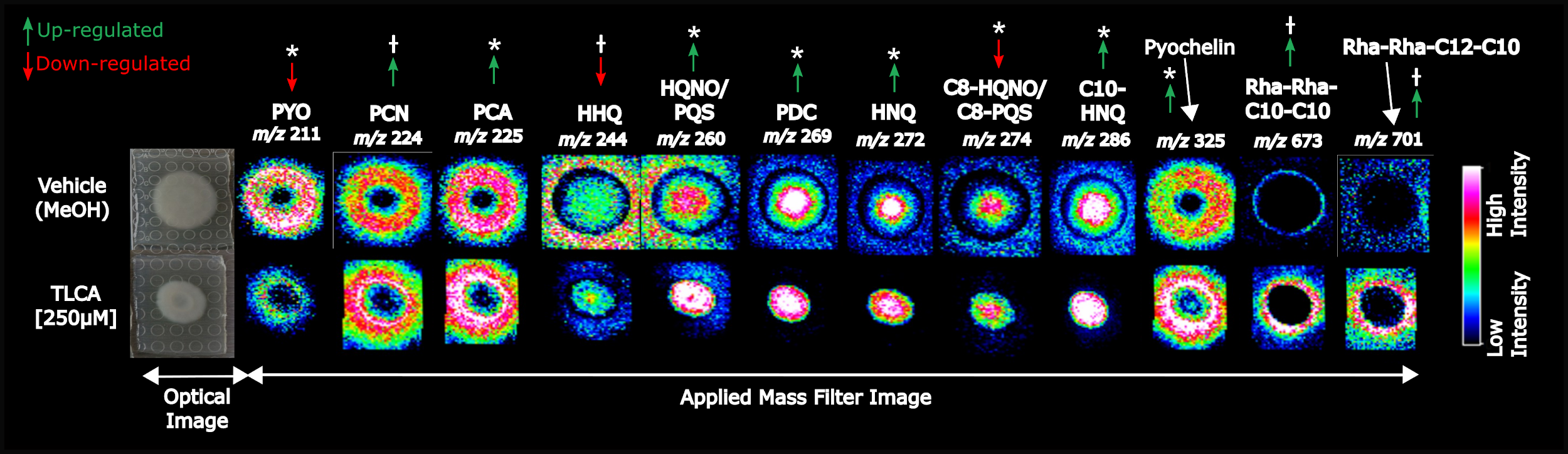

**Figure S4**: Replicate two IMS analysis of the twelve characterized metabolites seen in Figure 2. Statistical analysis reveals where a signal has altered regulation within the colony (*) or in the surrounding agar (ᵻ) (p<0.05).

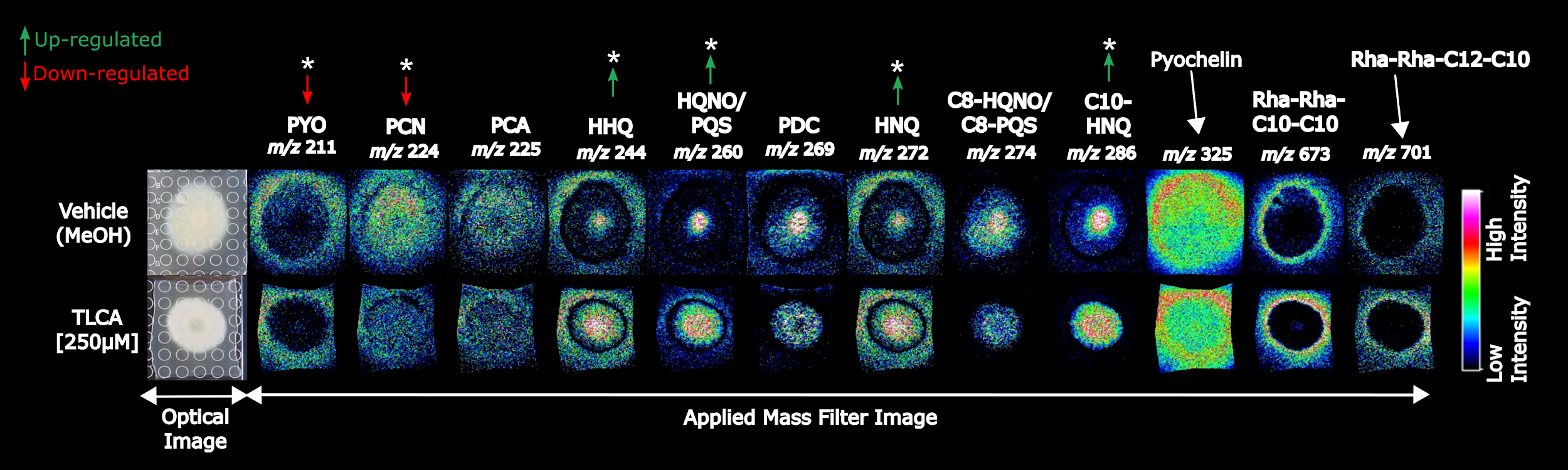

**Figure S5**: Replicate three IMS analysis of the twelve characterized metabolites seen in Figure 2. Statistical analysis reveals where a signal has altered regulation within the colony (*) or in the surrounding agar (ᵻ) (p<0.05).

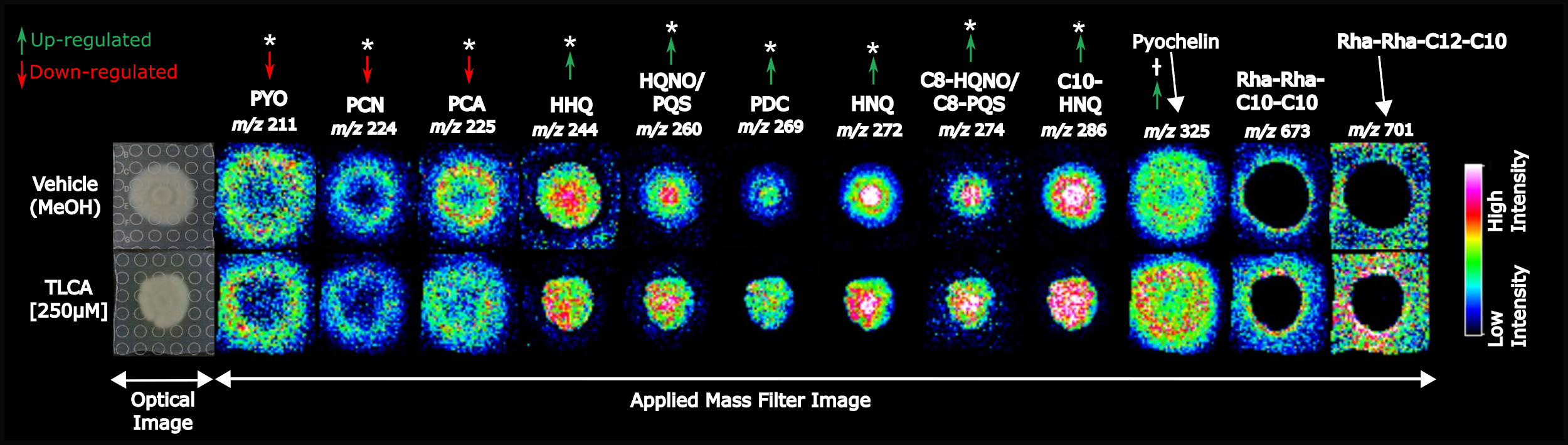

**Figure S6:** IMS of PA14 mutants revealed TLCA exposure induced an increase in pyochelin production in only the *pqsH* knockout, resembling the trend observed in wild type experiments. ♱♱ denotes the observed regulation was statistically significant over three biological replicates (p<0.05)

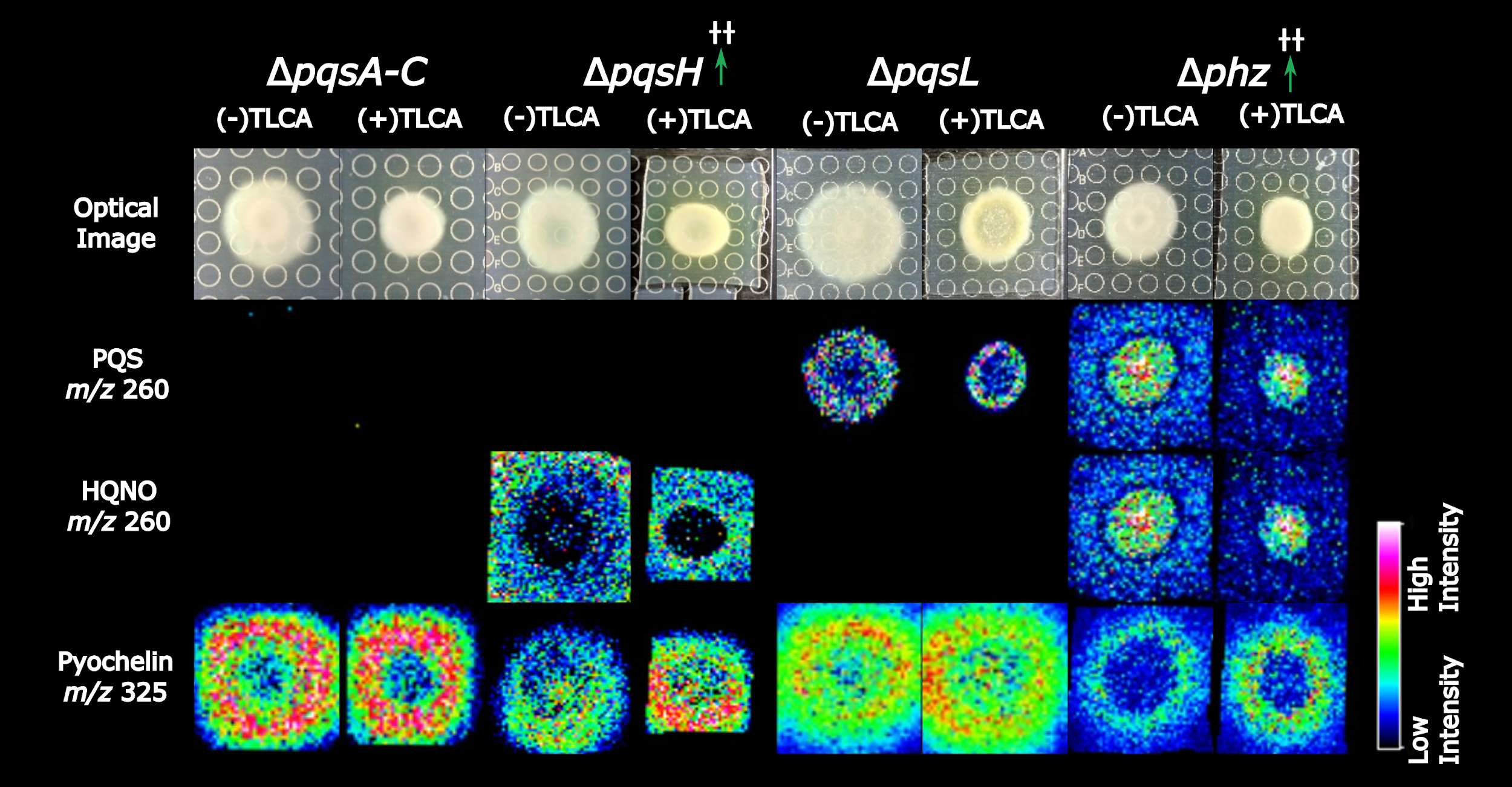

**Figure S7:** HPLC chromatogram and UV-Vis spectra of phenazine-1-carboxylic acid (PCA) standard (green solid line) and PA14 extract (red dashed line). PCA is characterized by UV maxima at 250 nm and 370 nm.

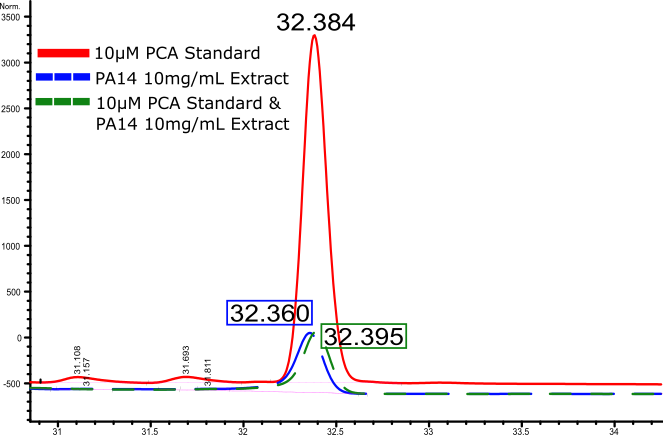

**Figure S8:** GNPS result comparing raw tandem MS/MS spectrum (black/top) to library hit for pyochelin (green/bottom) confirming the identification of pyochelin in the PA14 extract with a cosine similarity score of 0.93.
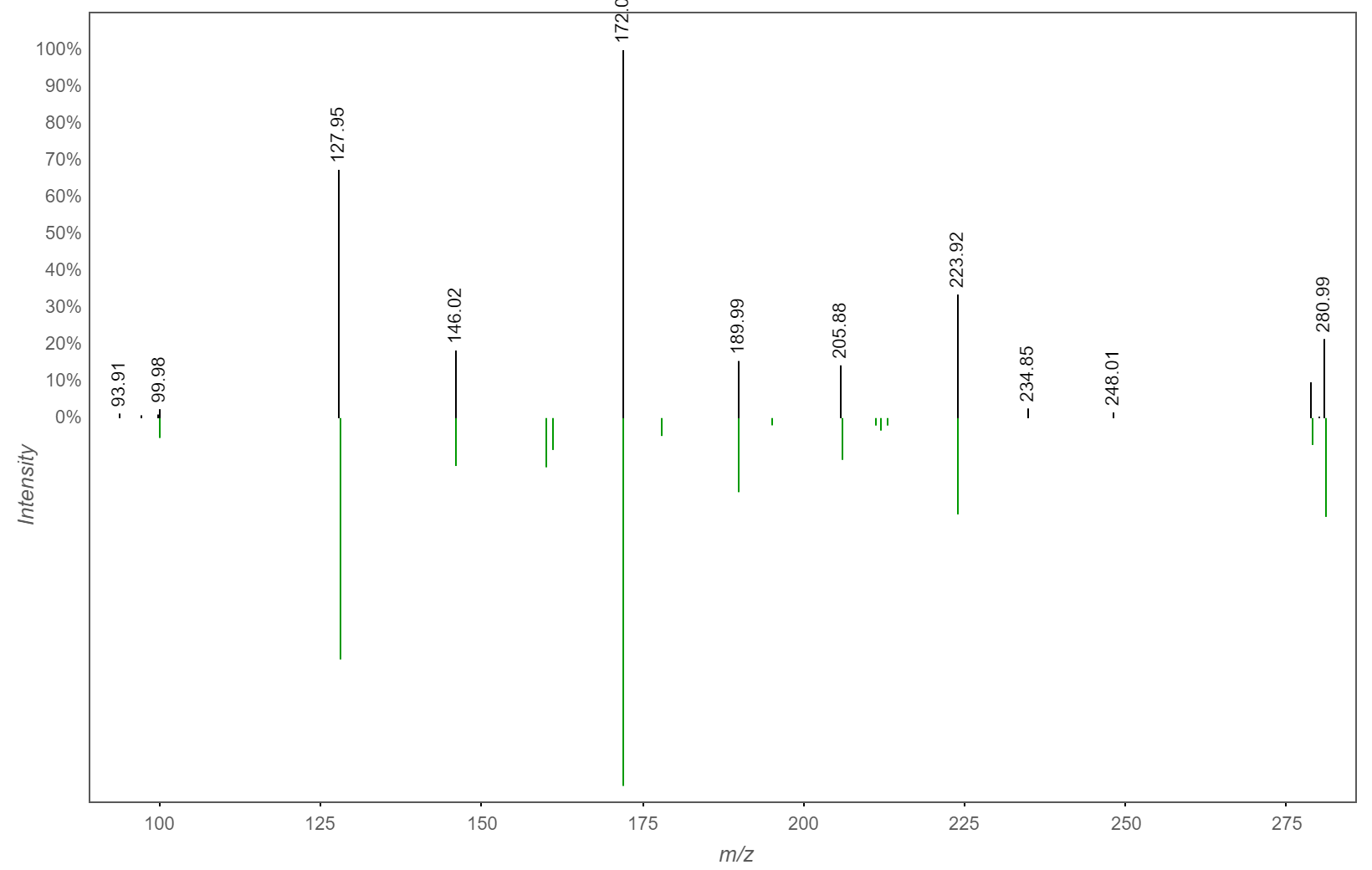

**Figure S9:** HPLC chromatogram and UV-Vis spectra of isolated pyochelin (PCH) from PA14 extract. Fe-bound PCH is characterized by UV maxima at 237 nm, 255 nm, and 325 nm [[3]](https://paperpile.com/c/xWBxHG/gKtN).

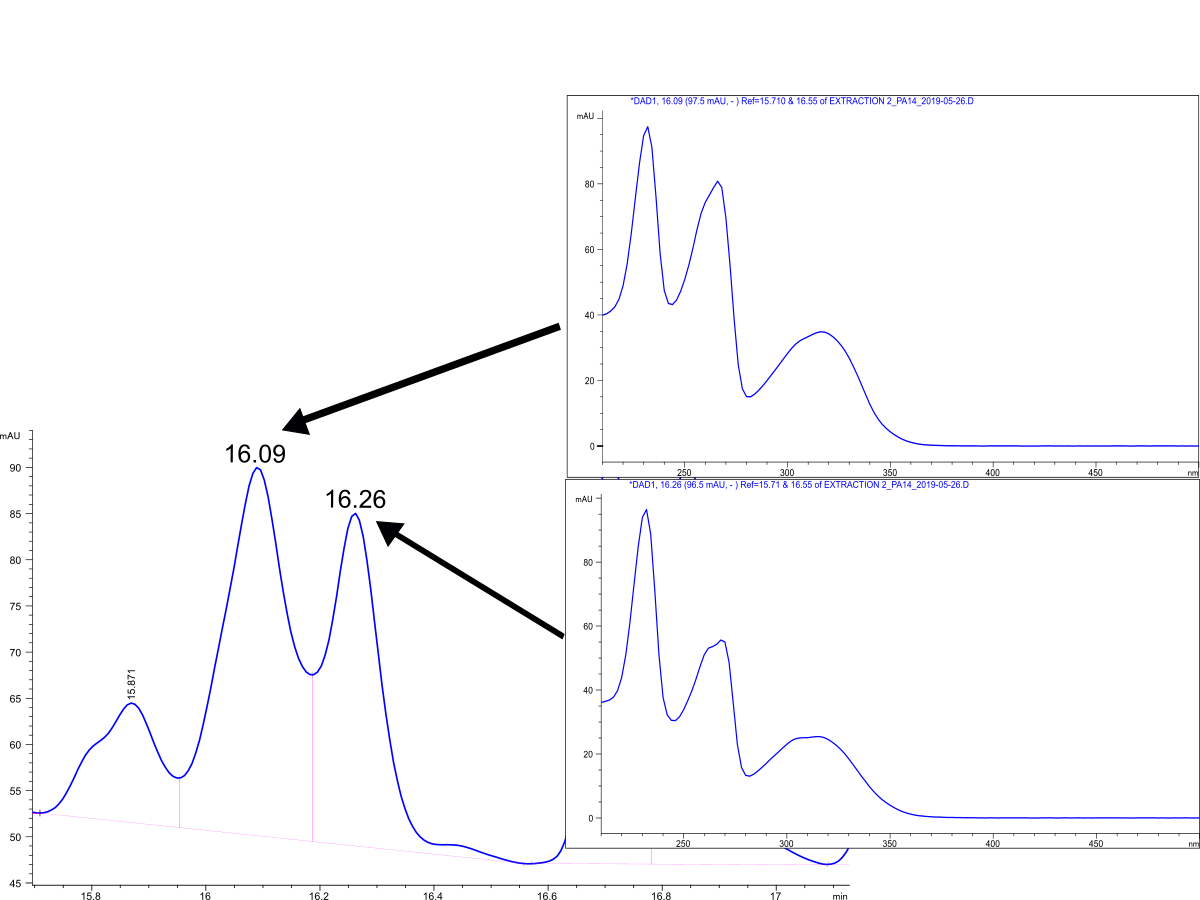

**Table S1**: Statistically significant signals from IMS experiments. Burker’s SCiLS software identified that all signals had significant altered regulation when exposed to TLCA whether within the bacterial colony (*) or from excretion in the surrounding agar (ᵻ) (p<0.05). NS = not significant.

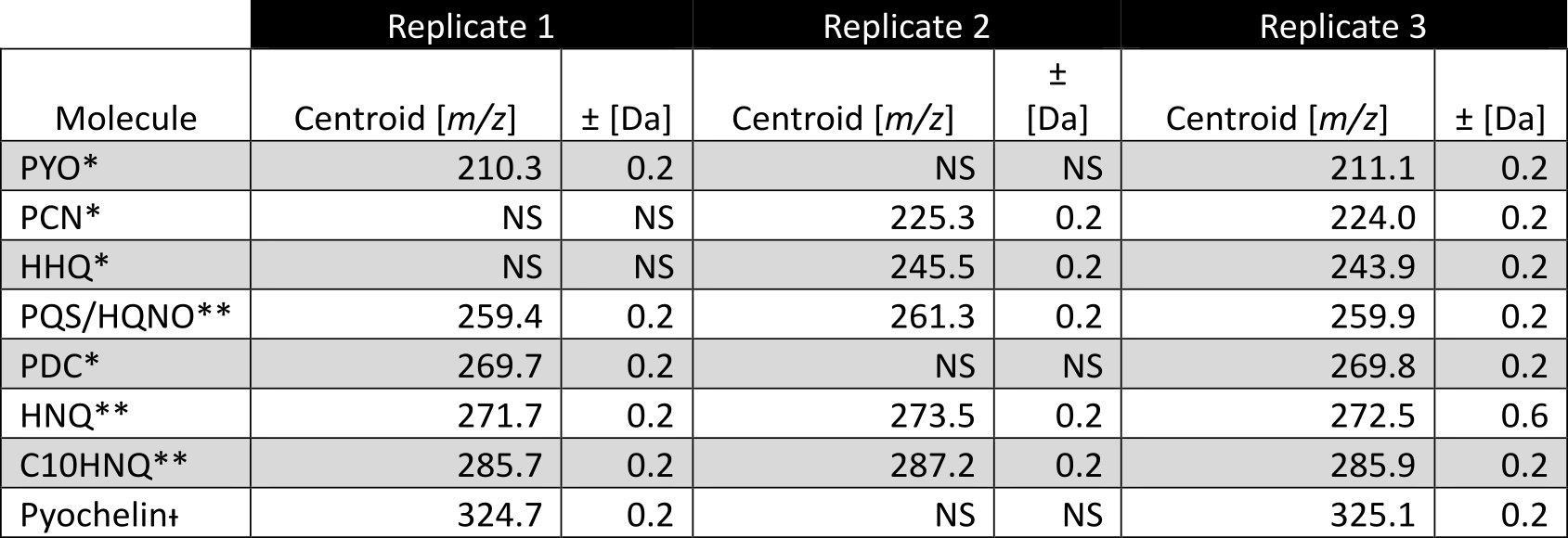

**Table S2**: Calculated Mass Error from IMS analysis. Using Matrix cluster as a reference the average mass error for the experiment was calculated.

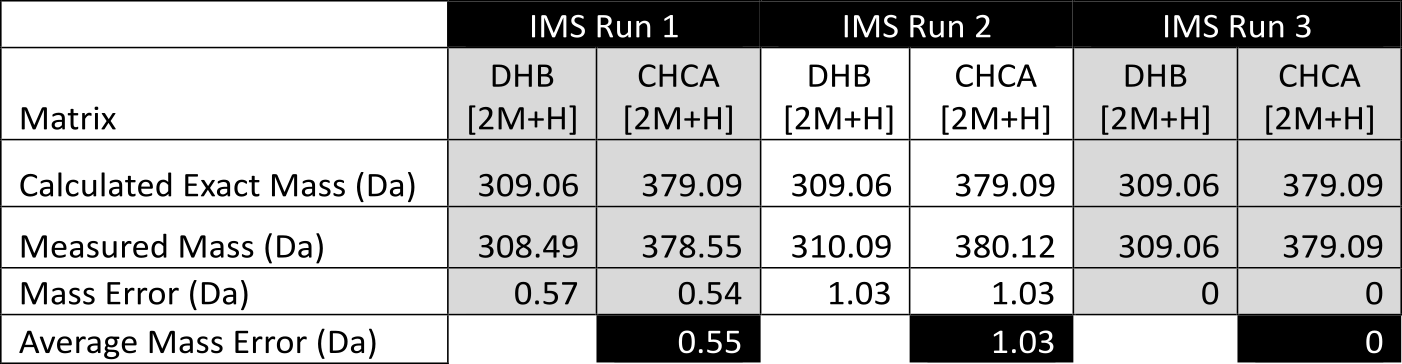

**Table S3**: Pairwise comparisons of the survival curves of all groups from wax moth virulence assay. Comparison is significant if p<0.05.

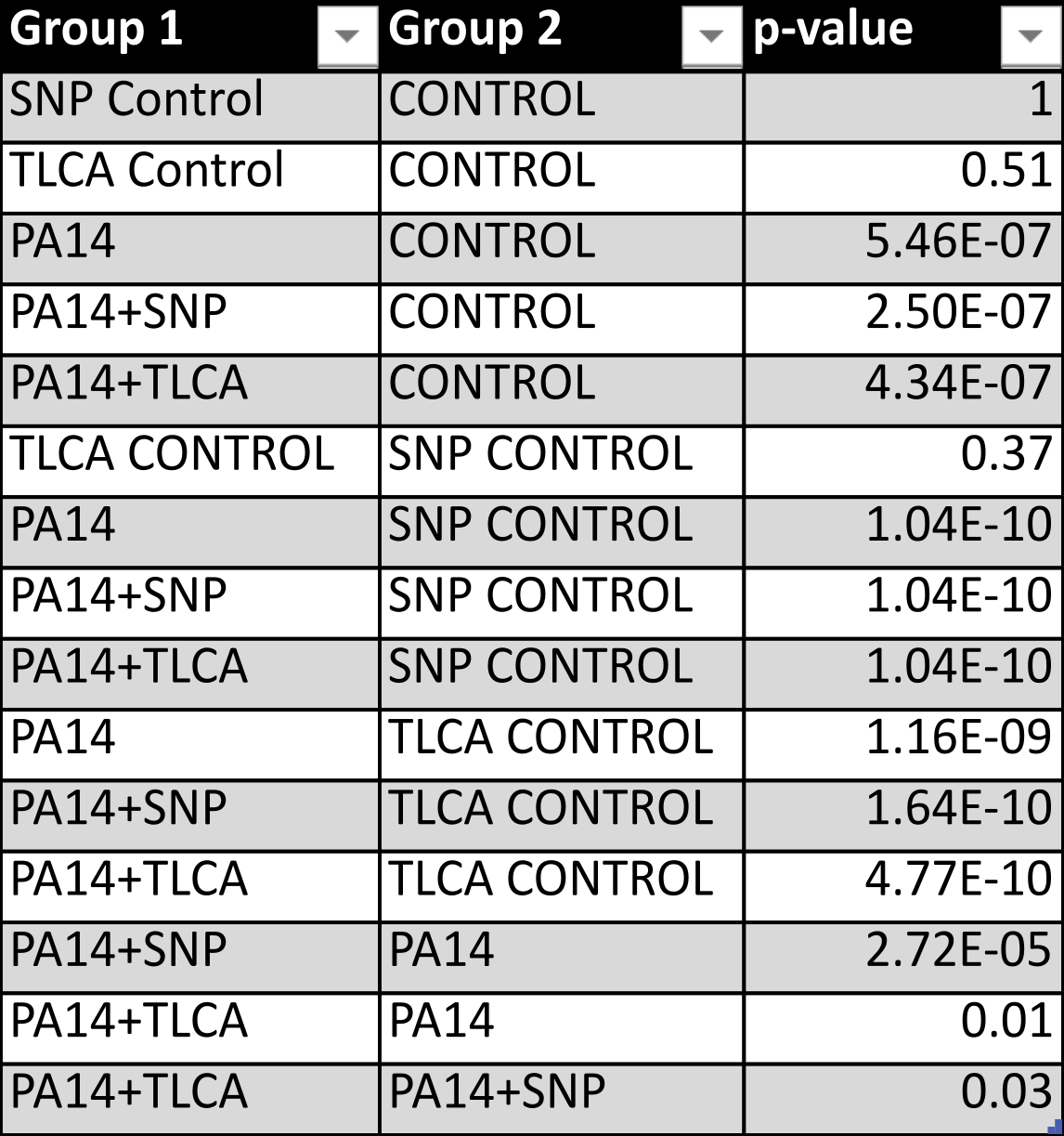

**Table S4**: Statistically significant signals of pyochelin from mutant PA14 IMS experiments (p<0.05). Burker’s SCiLS software identified when pyochelin had significant altered regulation when exposed to TLCA; upregulated (green), downregulated (red), or NS = not significant.

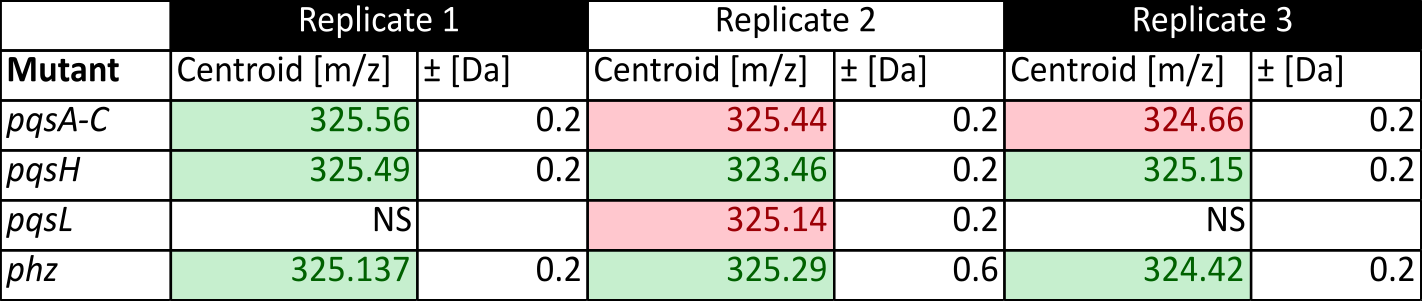

**Table S5**: Mass errors from *pqsA-C* replicate IMS experiments.

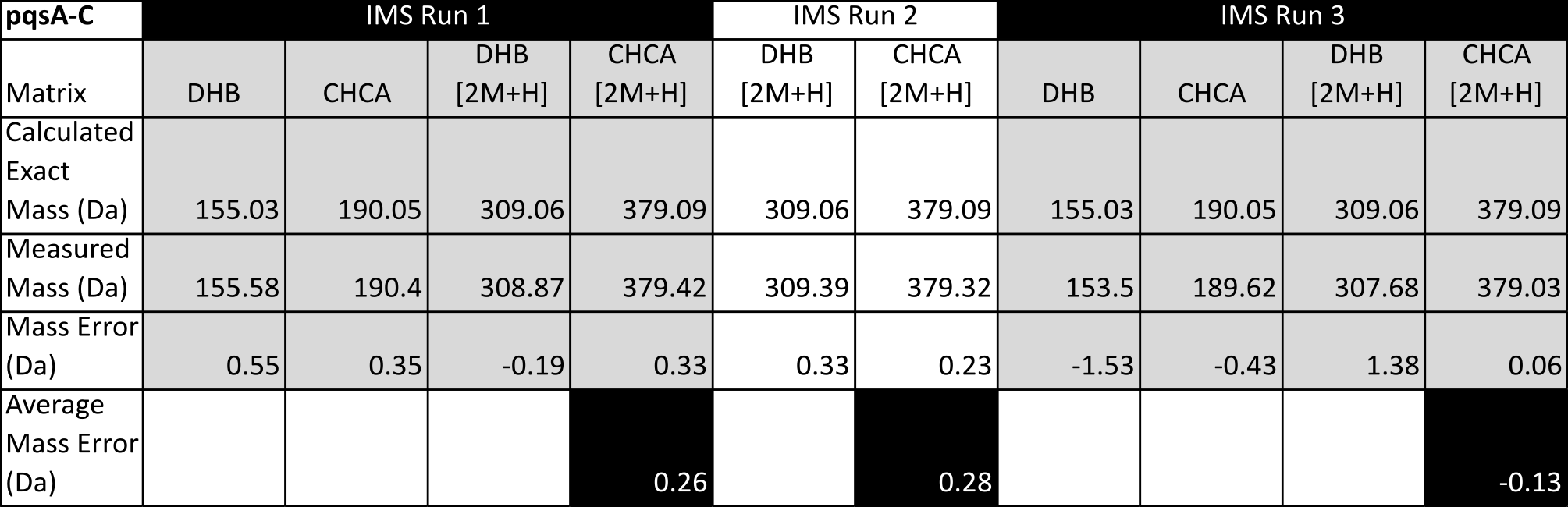

**Table S6**: Mass errors from *pqsH* replicate IMS experiments.

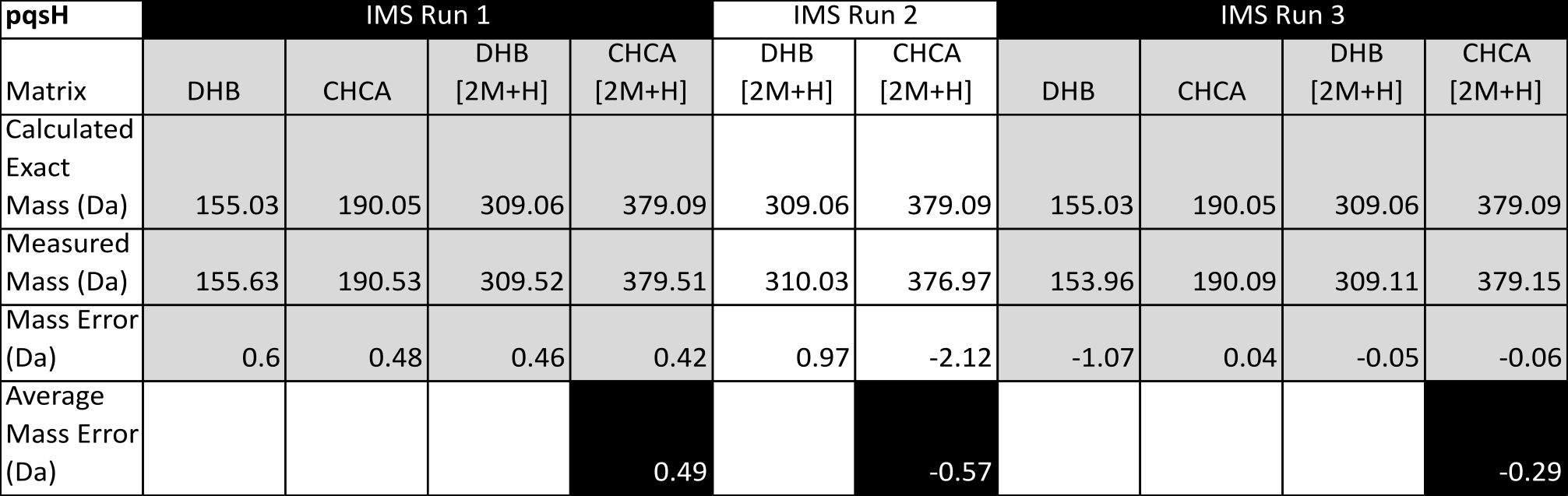

**Table S7**: Mass errors from *pqsL* replicate IMS experiments.

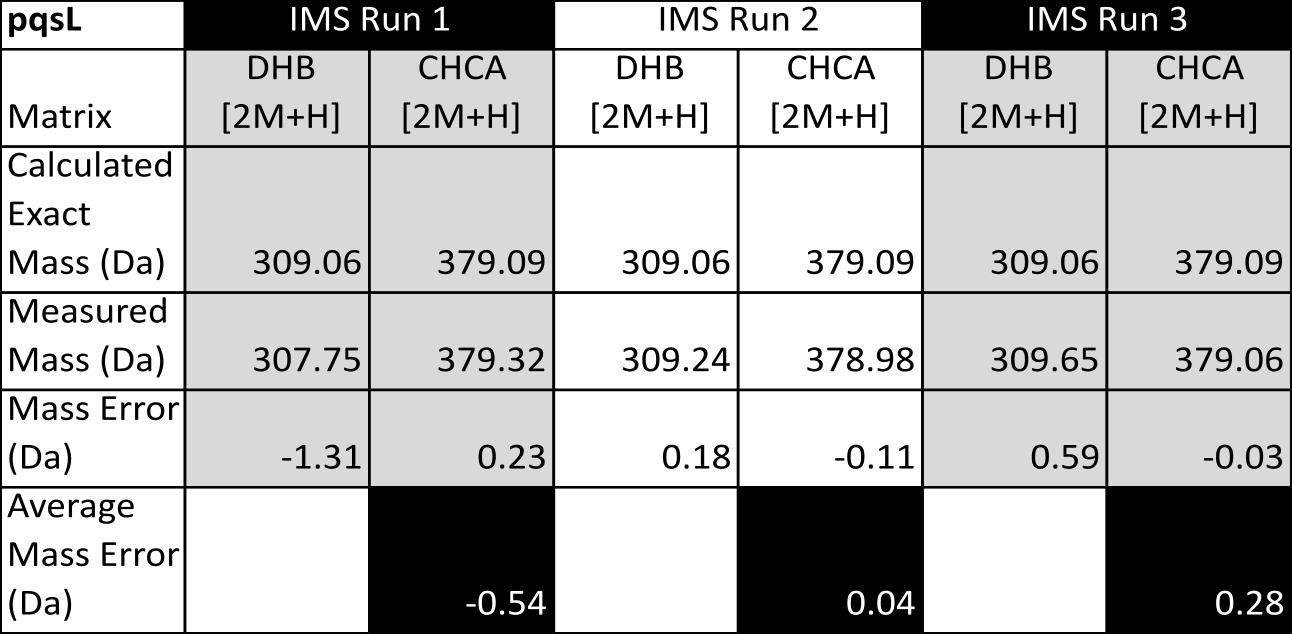

**Table S8**: Mass errors from *phz* replicate IMS experiments.

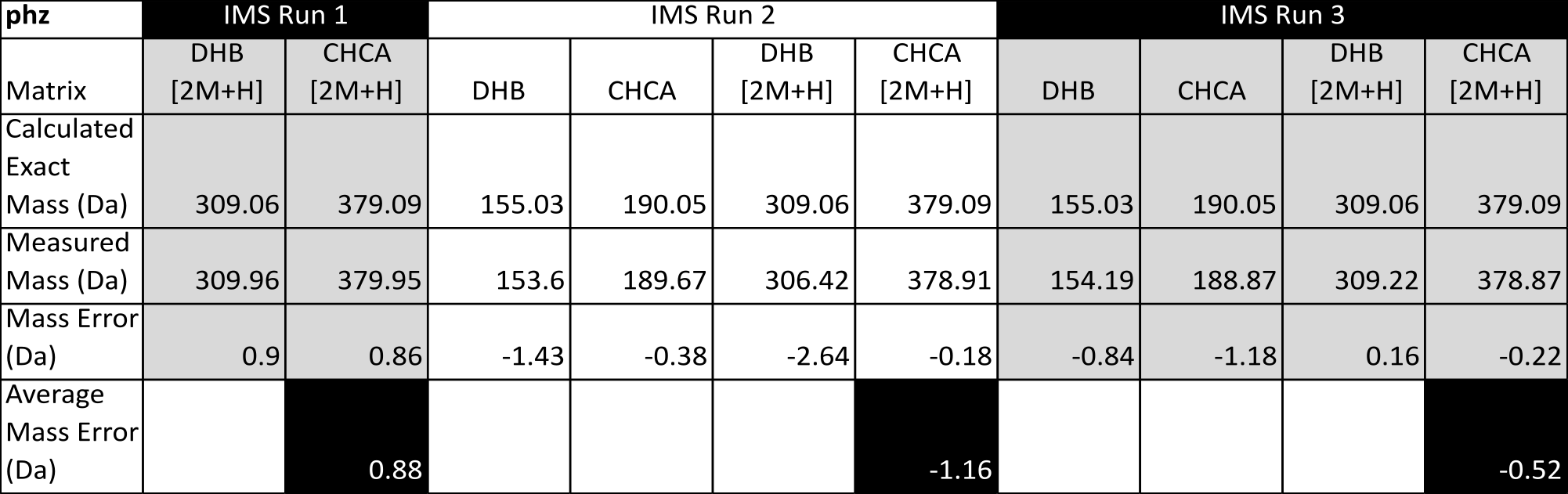

**Table S9**: Complete fold change analysis of PCA and PCH in PA14 strains treated with TLCA.

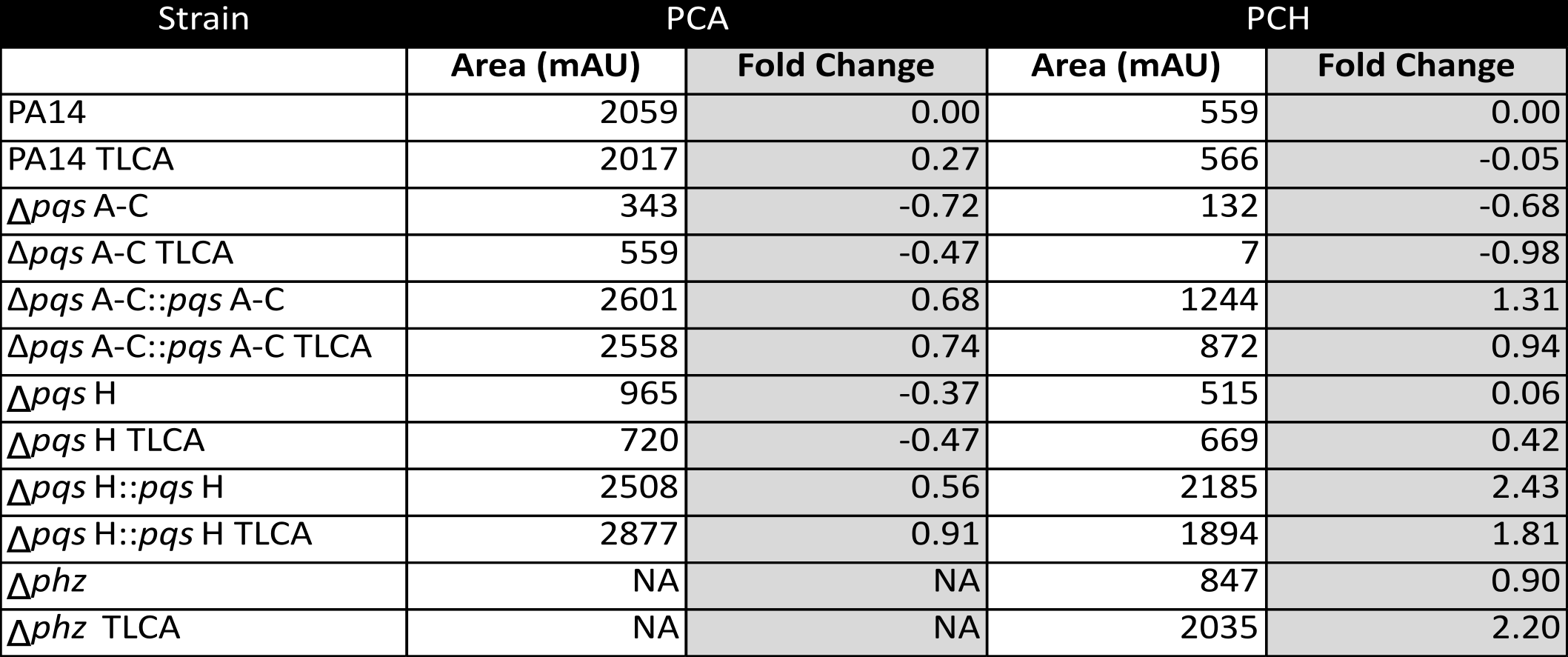

**Table S10.** Strains, plasmids and primers utilized in this study.

| **Strain** | **Number** | **Relevant characteristics** | **Source** |
| --- | --- | --- | --- |
| ***Pseudomonas aeruginosa*** | | | |
| UCBPP-PA14 | LD0 | Clinical isolate UCBPP-PA14 | [[4]](https://paperpile.com/c/xWBxHG/hQ74) |
| Clinical isolate UCBPP-PA14 | LMS1 | Clinical isolate UCBPP-PA14 | M. Banzhaf EMBL |
| PA14 ∆*phz1/2* | LD24 | PA14 with deletions in *phzA1-G1* and *phzA2-G2* operons | [[5]](https://paperpile.com/c/xWBxHG/p4RQ) |
| PA14 ∆*pqsA-C* | LD732 | PA14 with deletions of *pqsAI* (PA14_51430), *pqsB* (PA14_51420) and *pqsCI* (PA14_51410) in the *pqsA-E* operon | [[6]](https://paperpile.com/c/xWBxHG/krdx) |
| PA14 ∆*pqsH* | LD685 | PA14 with deletions of *pqsH* (PA14_30630) | [[6]](https://paperpile.com/c/xWBxHG/krdx) |
| PA14 ∆*pqsL* | LD701 | PA14 with deletions of *pqsL* (PA14_09700) | [[6]](https://paperpile.com/c/xWBxHG/krdx) |
| PA14 ∆*pqsA-C*::*pqsA-C* complement | LD3211 | LD732 parent with deleted region replaced with wildtype sequence | this study |
| PA14 ∆*pqsH*::*pqsH* complement | LD3212 | LD685 parent with deleted region replaced with wildtype sequence | this study |
| PA14 ∆*pqsL*::*pqsL* complement | LD3213 | LD701 parent with deleted region replaced with wildtype sequence | this study |
| ***Escherichia coli*** | | | |
| UQ950 | LD44 | *E. coli* DH5 λpir strain for cloning. F-∆(argF-lac)169φ80 dlacZ58(∆M15) glnV44(AS) rfbD1 gyrA96(NaIR ) recA1 endA1 spoT thi-1 hsdR17 deoR λpir+ | D. Lies,  Caltech |
| BW29427 | LD661 | Donor strain for conjugation. thrB1004 pro thi rpsL hsdS lacZ DM15RP4-1360 D(araBAD)567 DdapA1314::[erm pir(wt)] | W. Metcalf,  University of Illinois |
| ***Saccharomyces cerevisiae*** | | | |
| InvSc1 | LD676 | Used for yeast gap repair cloning | Invitrogen |

**Plasmids used in this study**

| **Plasmids** | **Sequence** | **Used to make plasmid number** |
| --- | --- | --- |
| pMQ30 | 7.5 kb mobilizable vector; oriT, sacB, GmR | [[7]](https://paperpile.com/c/xWBxHG/fVTy) |
| pLD3214 | *pqsA-C* PCR fragment introduced into pMQ30 by gap repair cloning in yeast strain InvSc1. | this study |
| pLD3215 | *pqsH* PCR fragment introduced into pMQ30 by gap repair cloning in yeast strain InvSc1. | this study |
| pLD3216 | *pqsL* PCR fragment introduced into pMQ30 by gap repair cloning in yeast strain InvSc1. | this study |

**Primers used in this study**

| **Primer number** | **Sequence** | **used to make plasmid number** |
| --- | --- | --- |
| LD1 | ggaattgtgagcggataacaatttcacacaggaaacagctAGAGGCTCCGATCACCCTAT | pLD3214 |
| LD4 | ccaggcaaattctgttttatcagaccgcttctgcgttctGAACCGTAGGTCAGGACCAG |  |
| LD9 | ggaattgtgagcggataacaatttcacacaggaaacagctCGCCTGTTCCTCAAGTACG | pLD3216 |
| LD12 | ccaggcaaattctgttttatcagaccgcttctgcgttCTCGAACAGGTGTTCCTCAATC |  |
| LD168 | ggaattgtgagcggataacaatttcacacaggaaacagctGATATCCACATCCACGGTGTC | pLD3215 |
| LD171 | ccaggcaaattctgttttatcagaccgcttctgcgttctgatCTTCTCGAAGATGCGGTAGC |  |

**Data availability**

GNPS Job Link 1:

<https://gnps.ucsd.edu/ProteoSAFe/status.jsp?task=3a542d7c1f3c4cd5b69bb99433065af3>

GNPS Job Link 2:

<https://gnps.ucsd.edu/ProteoSAFe/status.jsp?task=b96ee4ae2f9640d3b3a2dcabeed08c3a>

LC-MS/MS raw data (in triplicate): (accession number: **MSV000083258**)

All IMS raw data (in triplicate): (accession number: **MSV000083413**)

[1. Wang M, Carver JJ, Phelan VV, Sanchez LM, Garg N, Peng Y, Nguyen DD, Watrous J, Kapono CA, Luzzatto-Knaan T, Porto C, Bouslimani A, Melnik AV, Meehan MJ, Liu W-T, Crüsemann M, Boudreau PD, Esquenazi E, Sandoval-Calderón M, Kersten RD, Pace LA, Quinn RA, Duncan KR, Hsu C-C, Floros DJ, Gavilan RG, Kleigrewe K, Northen T, Dutton RJ, Parrot D, Carlson EE, Aigle B, Michelsen CF, Jelsbak L, Sohlenkamp C, Pevzner P, Edlund A, McLean J, Piel J, Murphy BT, Gerwick L, Liaw C-C, Yang Y-L, Humpf H-U, Maansson M, Keyzers RA, Sims AC, Johnson AR, Sidebottom AM, Sedio BE, Klitgaard A, Larson CB, P CAB, Torres-Mendoza D, Gonzalez DJ, Silva DB, Marques LM, Demarque DP, Pociute E, O’Neill EC, Briand E, Helfrich EJN, Granatosky EA, Glukhov E, Ryffel F, Houson H, Mohimani H, Kharbush JJ, Zeng Y, Vorholt JA, Kurita KL, Charusanti P, McPhail KL, Nielsen KF, Vuong L, Elfeki M, Traxler MF, Engene N, Koyama N, Vining OB, Baric R, Silva RR, Mascuch SJ, Tomasi S, Jenkins S, Macherla V, Hoffman T, Agarwal V, Williams PG, Dai J, Neupane R, Gurr J, Rodríguez AMC, Lamsa A, Zhang C, Dorrestein K, Duggan BM, Almaliti J, Allard P-M, Phapale P, Nothias L-F, Alexandrov T, Litaudon M, Wolfender J-L, Kyle JE, Metz TO, Peryea T, Nguyen D-T, VanLeer D, Shinn P, Jadhav A, Müller R, Waters KM, Shi W, Liu X, Zhang L, Knight R, Jensen PR, Palsson BO, Pogliano K, Linington RG, Gutiérrez M, Lopes NP, Gerwick WH, Moore BS, Dorrestein PC, Bandeira N (2016) Sharing and community curation of mass spectrometry data with Global Natural Products Social Molecular Networking. *Nature biotechnology*, 34(8):828–837.](http://paperpile.com/b/xWBxHG/YJaj)
